# Single Cell Multimodal Analyses Reveal Epigenomic and Transcriptomic Basis for Birth Defects in Maternal Diabetes

**DOI:** 10.1101/2022.07.25.501463

**Authors:** Tomohiro Nishino, Sanjeev S. Ranade, Angelo Pelonero, Benjamin J. van Soldt, Lin Ye, Michael Alexanian, Frances Koback, Yu Huang, Nandhini Sadagopan, Arun Padmanabhan, Reuben Thomas, Joke G. van Bemmel, Casey A. Gifford, Mauro W. Costa, Deepak Srivastava

**Affiliations:** Gladstone Institutes, San Francisco, CA, USA; Roddenberry Center for Stem Cell Biology and Medicine at Gladstone, San Francisco, CA, USA; Division of Cardiology, Department of Medicine, University of California, San Francisco, CA, USA; Division of Cardiology, Department of Pediatrics, University of California, San Francisco, CA, USA; Department of Biochemistry and Biophysics, University of California, San Francisco, CA, USA

## Abstract

Birth defects occur in ∼6% of all live births and can be caused by combinations of genetic and environmental influences^1^. Large-scale DNA sequencing efforts are revealing genetic influences^2,3^, but investigations into the contributions of environmental factors have largely been limited to association studies with limited mechanistic insight. Hyperglycemia present in pre-gestational diabetic mothers is among the most frequent environmental contributor to congenital defects and results in an increased incidence of congenital heart defects and craniofacial anomalies^4^. However, the cell types involved and underlying mechanisms by which maternal hyperglycemia affects these regions are unknown. Here, we utilized multi-modal single cell analyses to reveal that maternal diabetes affects the epigenomic and transcriptomic state of specific subsets of cardiac and craniofacial progenitors during embryogenesis. A previously unrecognized subpopulation of anterior heart field progenitors expressing *Alx3* acquired a more posterior identity in response to maternal hyperglycemia, based on gene expression and chromatin status. Similarly, a sub-population of neural crest-derived cells in the second pharyngeal arch, which contributes to craniofacial structures, also displayed abnormalities in cell specification and patterning. Analysis of differentially accessible chromatin regions demonstrated that disrupted patterning was associated with increased intrinsic retinoic acid signaling in affected cell types in response to maternal diabetes and hyperglycemia. This work demonstrates how an environmental insult can have highly selective epigenomic consequences on discrete cell types leading to developmental patterning defects.

---

Cardiac and craniofacial birth defects often occur in conjunction due to shared cardio-pharyngeal and neural crest-derived progenitors, and reciprocal signaling between the two populations^5,6^. Numerous genetic syndromes affect both regions, as do environmental influences. Among the latter, pregestational maternal diabetes (PGDM), and the associated hyperglycemia, is relatively common given the increasing prevalence of type II diabetes worldwide, and leads to a 4 to 5-fold increase in the incidence of birth defects^7^. Similarly, heart and craniofacial regions are also severely affected upon rare external exposure to retinoic acid^8^. Retinoic acid embryopathy is related to disruption of anterior-posterior patterning, particularly through dysregulation of the Hox code in pharyngeal arches and specification of the anterior and posterior regions of second heart field progenitors that contribute to the outflow and inflow tracts of the heart, respectively^9,10^. The mechanisms by which hyperglycemia increases the risk for birth defects are unclear, although there is some evidence that oxidative stress and increases in the metabolite beta hydroxybutyrate may lead to epigenomic changes^11-14^. To address this question, we analyzed the epigenomic landscape during mouse development at single cell resolution and deciphered cell-specific mechanisms that result in malformations.

## Cardiac malformations in mouse maternal diabetes mellitus

To investigate the cellular consequences of PGDM during embryonic cardio-pharyngeal development, we induced diabetes mellitus (DM) in female mice by intraperitoneal delivery of streptozotocin (STZ) for five consecutive days, resulting in loss of insulin-producing cells. Females with blood glucose levels greater than 250 mg/dL were mated with untreated males (mean blood glucose levels in vehicle (VEH): 180.1 ± 4.0 mg/dl, in STZ: 356.4 ± 13.2 mg/dl) (**Fig. 1a** and **Extended Data Fig. 1a**). As expected, embryos from PGDM females had significantly higher frequency of cardiac malformations that included ventricular septal defects, atrial septal defects, and right ventricular hypoplasia when compared to embryos from VEH treated females (**Extended Data Fig. 1b** and **Extended Data Table 1**). Neural tube defects and craniofacial defects also occurred in the STZ-induced PGDM model as previously reported, mimicking the human condition^15-17^.

**Fig. 1.**
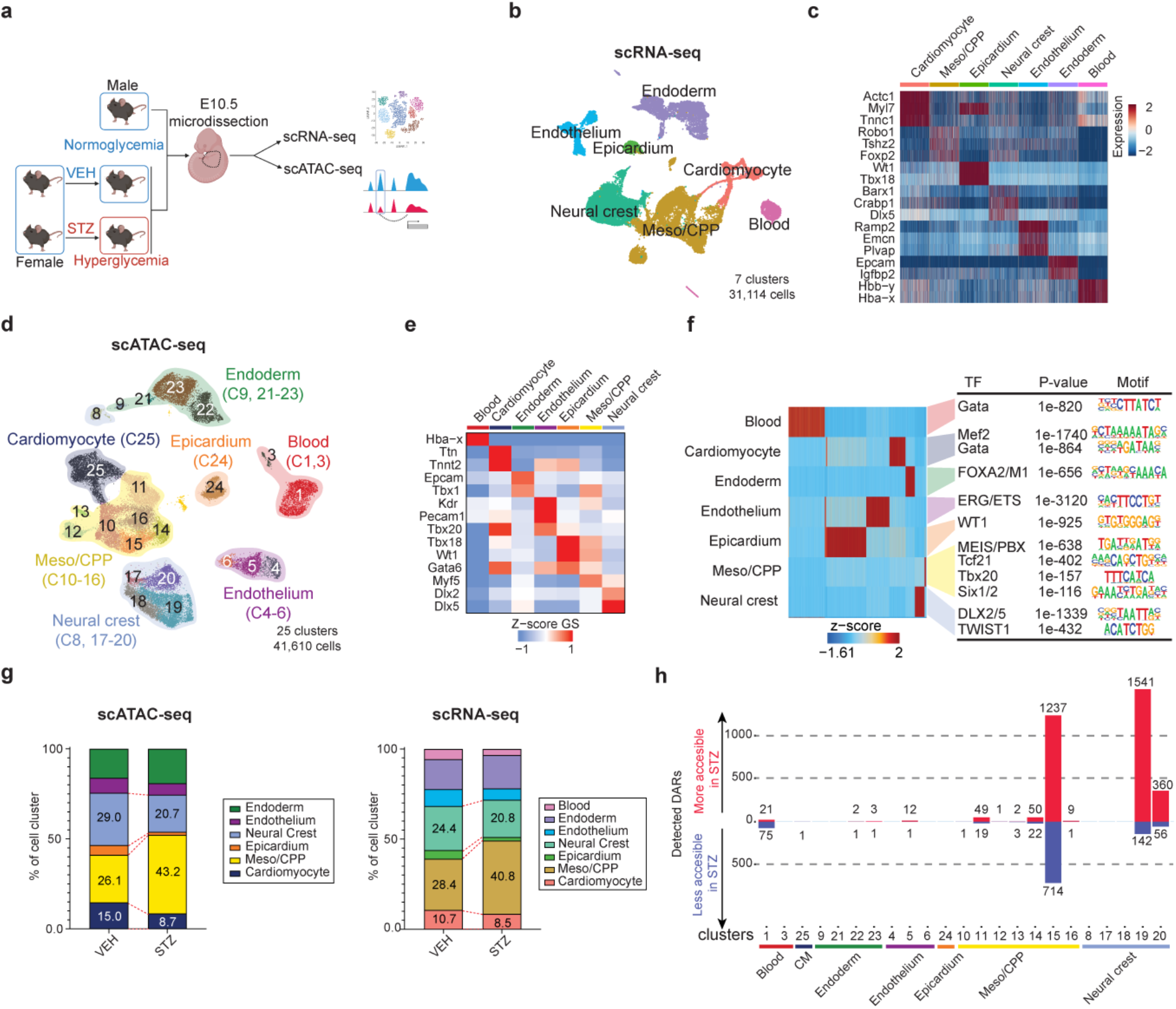
Integrated Analysis of scRNA-seq and scATAC-seq Reveals Highly Selective Chromatin Accessibility Alterations in PGDM. **a**, Strategy for the *in vivo* pregestational diabetes mellitus (PGDM) model after treatment of either vehicle (VEH) or streptozotocin (STZ). Samples were collected from area indicated with dotted line at embryonic day 10.5 (E10.5) and split to perform scRNA-seq and scATAC-seq using the 10x Genomics Chromium system. **b**, Uniform manifold approximation and projection (UMAP) presentation of scRNA-seq of all captured cells from the cardio-pharyngeal area at E10.5 colored by cluster identity. Cells were collected from three E10.5 embryos from VEH treated females and three from STZ treated females. **c**, Expression heat map of marker genes of each defined populations. Scale indicates z-scored expression values. All genes represented have an adjusted p-value less than 1 × 10^−10^ (Wilcoxon rank-sum test). **d**, UMAP representation of all captured cells in scATAC-seq colored by cluster identity and grouped based on cell types. Cells were collected from the same embryos as those used for scRNA-seq shown in b. **e**, Heatmap of Gene Scores (GS) from curated marker genes based on existing knowledge and scRNA-seq data for each cell type. Scale indicates z-scored GS values. **f**, Top enriched transcription factor (TF) motifs in detected marker peaks (FDR < 1.00e-08 and Log2FC > 2) for each cell type analyzed using HOMER *de novo* analysis. **g**, Population distribution of each cell type normalized to total number of cells per sample in scATAC-seq (left) and scRNA-seq (right). Statistics performed by permutation test in scATAC-seq data (STZ vs. VEH), for Meso/CPP, FDR<0.001, Log2FD=0.79; for Cardiomyocyte, FDR<0.001, Log2FD=-0.71; for Neural crest, FDR<0.001, Log2FD=-0.43; for Endoderm, FDR<0.001, Log2FD=0.32; for Endothelium, FDR<0.001, Log2FD=-0.30; for Epicardium, FDR<0.001, Log2FD=-1.59. Statistics performed by permutation test in scRNA-seq data (STZ vs. VEH), for Meso/CPP, FDR<0.001, Log2FD=0.54; for Cardiomyocyte, FDR<0.001, Log2FD=-0.36; for Neural crest, FDR<0.001, Log2FD=-0.25; for Endoderm, FDR<0.001, Log2FD=0.15; for Endothelium, FDR<0.001, Log2FD=-0.62; for Epicardium, FDR<0.001, Log2FD=-1.38; for Blood, FDR<0.001, Log2FD=-0.65. **h**, Distribution of detected differentially accessible regions (DARs) between VEH and STZ groups in each sub cluster across all cell type. (FDR < 0.05 and |LogFC| > 1). Meso/CPP; Mesodermal cardio-pharyngeal progenitors

## Cell population changes in cardio-pharyngeal development due to PGDM

To determine how PGDM affects the transcriptional and epigenomic state within individual cells during cardio-pharyngeal development, we conducted single cell RNA-seq (scRNA-seq) and single-cell sequencing assay for transposase-accessible chromatin (scATAC-seq) on the cardiac and surrounding pharyngeal region at E10.5 from VEH- or STZ-treated females (**Fig. 1a** and **Extended Data Fig. 2a**). At this time point, the precursors of the major areas affected in this model are present, including neural crest-derived cells. After dissociating the dissected cardio-pharyngeal region into single cells, each individual sample was split for parallel scRNA-seq and scATAC-seq experiments. Using unsupervised clustering with Seurat^18^, single-cell transcriptomes of over 31,000 cells were clustered, identified and labeled based on known marker genes. This initial analysis confirmed the presence of all expected cell types at E10.5 (**Fig. 1b, c, Extended Data Fig. 2b, left, c**).

In parallel, analysis of single-cell chromatin accessibility data by scATAC-seq of 41,610 high-quality cells with batch correction identified 25 clusters using ArchR^19^. Cluster identity was assigned based on gene activity score of known cell type specific marker genes, a measure of the chromatin accessibility of the promoter region and gene body of each gene (**Fig. 1d, e, Extended Data Fig. 2b, right**). The specific peaks for each cluster were then identified, and transcription factor (TF) motifs enriched in those peaks were analyzed by HOMER^20^, representing distinct cell types (**Fig. 1f**). Furthermore, integration of scRNA-seq and scATAC-seq data supported our cluster annotations based on chromatin accessibility, demonstrating that all cell types expected from our microdissection were captured in the scATAC-seq experiment (**Extended Data Fig. 2d**).

Annotation of scATAC-seq data suggested that PGDM affected the cellular population distribution, leading to an increase in cardio-pharyngeal mesodermal progenitors (26.1% (VEH) vs. 43.2% (STZ), FDR<0.05), while decreasing more differentiated cardiomyocytes (15.0% (VEH) vs. 8.7% (STZ), FDR<0.05) and neural crest derivatives (29.0% (VEH) vs. 20.7% (STZ), FDR<0.05) (**Fig. 1g, left)**. scRNA-seq data corroborated an increase in cardio-pharyngeal mesodermal progenitors with decreases in neural crest cells and cardiomyocytes (**Fig. 1g, right**). Interestingly, all of the affected cell populations are directly involved in the types of cardiac and craniofacial defects observed in the setting of PGDM.

## Cell type specific epigenomic and transcriptional consequences of PGDM at cellular resolution

While numbers of cells may be important, dysregulation of gene networks within specific cell types may reveal the mechanistic basis for the morphogenetic defects associated with PGDM. We therefore performed peak calling for each scATAC-seq cluster, identifying a total of 492,330 sites, with about 76% being distal peaks (> 3 kb away from transcription start site (TSS)) (**Extended Data Fig. 2e, f**). Differentially accessible chromatin regions (DARs) were analyzed between the control (VEH) and hyperglycemic (STZ) conditions for each individual cluster. Surprisingly, of the total 4324 DARs detected, ∼97% were concentrated in only two cell types: mesodermally-derived cardio-pharyngeal progenitors (48.8%) and neural crest-derived cells (48.5%). Upon deeper analysis of these two cell compartments, we found that DARs were mostly enriched in a subset of clusters within each cell type. Among the seven subclusters of cardio-pharyngeal mesodermal progenitors, over 90% of DARs were detected in a single cluster (C15). Similarly, of the five neural crest sub-clusters, only two had DARs, with 80% of those being in a single cluster (C19) (**Fig. 1h**). We did not observe any correlation between detected DAR numbers and cell number or cellular distribution between conditions in each cluster. These results indicate that maternal PGDM-induced changes in embryonic chromatin accessibility are exquisitely cell-specific despite the universal exposure to hyperglycemia, and the cell types most affected in our data are consistent with those most critical for PGDM-related congenital diseases.

Given the concentration of DARs in neural crest-derived cells and cardio-pharyngeal mesodermal cells, we re-clustered the scRNA-seq data, focusing on those populations. Neural crest cell sub-clusters were annotated based on expression of *Hox* genes and other markers of neural crest-derivatives that distinguish cells that reside in different pharyngeal and aortic arches (**Fig. 2a-c**)^21,22^. Six distinct neural crest-related clusters were identified representing pharyngeal arch 2 (PA2), PA3, PA4/6, migrating neural crest progenitors (NC-prog), and the neural crest-derived smooth muscle cell progenitors (SMC-prog) and smooth muscle cells (SMC) of the aortic arch arteries and outflow tract. Analyses of the cellular distribution among each sample showed that PGDM led to an increase of undifferentiated NC-prog (2.7% (VEH) vs. 9.5% (STZ), FDR<0.05) with a concomitant decrease in the Rgs5-positive SMC progenitor population of the cardiac outflow tract (22.7% (VEH) vs. 17.8% (STZ), FDR<0.05) (**Extended Data Fig. 3a**). Integration of the annotated neural crest scRNA-seq data with the neural crest portion of the scATAC-seq data revealed that the neural crest-related clusters with the largest number of DARs, C19 and C20, represented cells from PA2 and PA3/4/6, respectively (**Fig. 2d-f, and Extended Data Fig. 3b**).

**Fig. 2.**
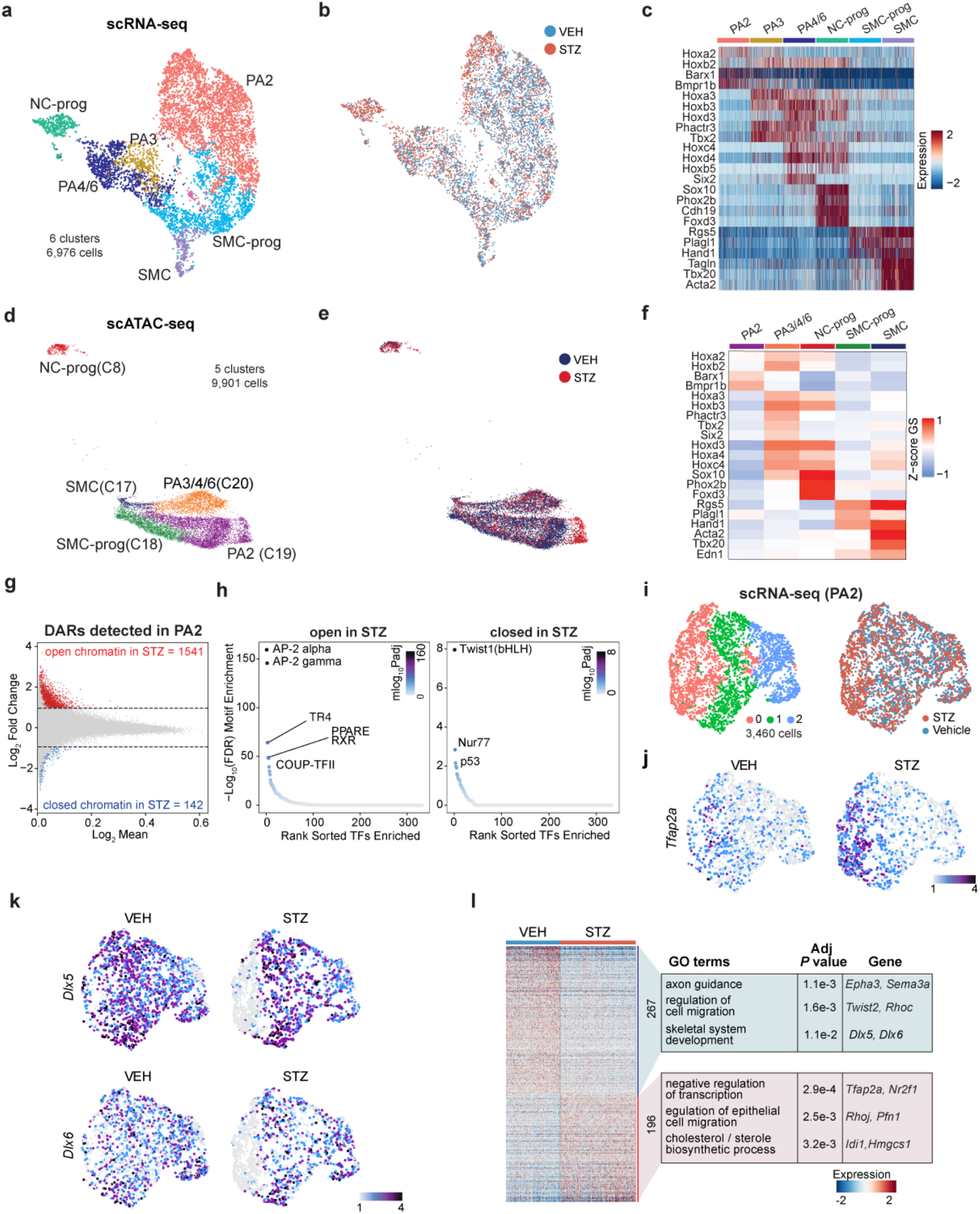
Maternal Diabetes Disrupts the Epigenomic Landscape of Craniofacial Neural Crest Cells in the Second Pharyngeal Arch. **a**, scRNA-seq UMAP representation of the neural crest cell population from Figure 1b, including VEH (n=3) and STZ cells (n=3), colored by cluster identity. **b**, scRNA-seq UMAP representation of neural crest cells colored by conditions. **c**, Heatmap for marker gene expression in the neural crest cell sub-populations. The Hox genes delineate spatial information. Scale indicates z-scored expression values. **d**, scATAC-seq UMAP representation of the neural crest cell population colored by cell type identity with original cluster numbers in parenthesis. **e**, scATAC-seq UMAP representation of the neural crest cell population colored by conditions. **f**, Heatmap of Gene Scores (GS) of curated marker genes based on scRNA-seq data for each neural crest sub-type. Scale indicates z-scored GS values. **g**, MA plot of differentially accessible regions (DARs) in PA2 population between VEH and STZ. Red dots represent the more accessible (open) chromatin region in STZ (FDR <= 0.05 & Log2FC >= 1) and blue dots represent less accessible (closed) chromatin region in STZ (FDR <= 0.05 & Log2FC <= -1). **h**, Enriched TF binding motifs in more accessible (left) or less accessible (right) DARs in STZ within PA2. **i**, scRNA-seq UMAP representation of PA2 neural crest cells. UMAP colored by cluster identity (left) or by conditions (right). **j**, Expression of *Tfap2a* on UMAP of PA2 neural crest cells in VEH or STZ embryos. Scale indicates z-scored expression values. **k**, Expression of *Dlx5* and *Dlx6* on UMAP of PA2 neural crest cells in VEH or STZ embryos. Scale indicates z-scored expression values. **l**, Heatmap representation of differentially expressed genes (DEGs) in PA2 subset cluster 0 between VEH and STZ (adjusted p-val < 0.05 and Log2FC > 0.25). Top GO terms enriched in upregulated or downregulated genes are shown with representative genes for each GO term. Scale indicates z-scored expression values. NC-prog, neural crest cell progenitors; PA2, pharyngeal arch 2; PA3, pharyngeal arch 3; PA4/6, pharyngeal arch 4/6; SMC, smooth muscle cells; SMC-prog, smooth muscle cell progenitors.

We performed TF motif analysis among the DARs in PA2 cranial neural crest cells and PA3/4/6 cardiac neural crest cells to determine how the gene regulatory regions were affected by PGDM and hyperglycemia. The AP2 motif, typical of undifferentiated neural crest cells^23,24^, was the most enriched in DARs with increased accessibility upon STZ exposure in both clusters, suggesting an impairment of differentiation in these populations. Retinoic acid-related RXR motifs were also enriched in the more accessible DARs of PA2, while the TWIST1 motif was enriched in less accessible DARs, supporting impairment of differentiation and consistent with *Twist1*’s role in cranial neural crest cell differentiation^25,26^ (**Fig. 2g, h and Extended Data Fig. 3c, d**). Similarly, in the PA3/4/6 cardiac neural crest, SIX, TBX, and SMAD2/4 motifs were enriched in DARs with reduced accessibility upon STZ treatment, suggesting the disruption of the core differentiation regulatory networks^27,28^ (**Extended Data Fig. 3d**). Although unsupervised clustering could not differentiate PA3 and PA4/6 in the C20 population, we were able to distinguish between PA3 and PA4/6 cells within C20 based on gene scores of several genes enriched in PA3 or PA4/6 by scRNA-seq, including *Hox* genes, *Six2*, and *Tbx2* (**Fig. 2d and Extended Data Fig. 3, f**). Subsequent DAR analysis between VEH and STZ in PA3 or PA4/6 showed an unbalanced difference in numbers of detected DARs, with changes detected almost exclusively in PA4/6, which is involved in pulmonary artery and aortic arch development (**Extended Data Fig. 3g**). TF motif enrichment analysis on DARs detected in PA4/6 reproduced the DAR findings in C20 (**Extended Data Fig. 3d**), including AP2 motif enrichment in more accessible loci in STZ and SMAD motif enrichment in less accessible loci in STZ, suggesting the chromatin accessibility change in C20 likely derived from PA4/6 changes (**Extended Data Fig. 3h**). In addition, the Sox TF motifs that form the core network in neural crest cell specification and migration^29^ were recognized in the more accessible DAR loci in STZ (**Extended Data Fig. 3h**). In summary, our results indicate that in the setting of PDGM/hyperglycemia, cells in PA2 and Six2-high PA4/6 display transcriptomic and epigenomic changes suggesting a less differentiated status.

Within the PA2 cells, a subpopulation was highly enriched in the STZ group compared to VEH group (**Fig. 2b, e**) leading us to investigate the identity and transcriptional consequences of PGDM in these cells. scRNA-seq reclustering of PA2 identified three distinct clusters, with most of the changes under STZ conditions localized to cluster 0 (**Fig. 2i**). This cluster was marked with *Tfap2a* expression, typical of the less differentiated neural crest cells, and was highly enriched upon STZ treatment, indicating that relatively undifferentiated cells within PA2 were most susceptible to PGDM and hyperglycemia, consistent with enrichment of the AP2 motif in more accessible DARs within PA2 (**Fig. 2h-j, and Extended Data Fig. 4a**). In addition, *Nr2f1*, which forms a TF complex with *Tfap2a* in cranial neural crest cells to maintain a relatively undifferentiated state^30^, was significantly upregulated in cluster 0 (**Extended Data Fig. 4b**).

*Tfap2a* is responsible for positional specification of neural crest cells along both the anterior-posterior (A-P) and dorsal-ventral (D-V) axis. The A-P axis is partly regulated through one of *Tfap2a’s* direct downstream target genes, *Hoxa2* ^31^, which was also significantly upregulated in cluster 0 among STZ offspring (**Extended Data Fig. 4c**). *Hoxa2* directly downregulates *Six2* and dictates A-P patterning^32^, which is consistent with the inverse correlation in expression levels of these two genes in PA2, PA3, and PA4/6, shown in Figure 2c. We also found *Dlx5* and *Dlx6*, which dictate D-V patterning in PA-related craniofacial development and are also downstream target genes of *Tfap2a* ^33^, were significantly downregulated in cluster 0 cells in STZ (**Fig. 2k and Extended Data Fig. 4d**).

In addition to pattern formation, the dysregulated genes are involved in cell differentiation and migration. For example, *Hoxa2* transiently activates *Meox1*, which is important for skeletal development from PA2 neural crest cells. While *Meox1* is expressed as early as E9.0, it is normally downregulated by E10.5 in PA2^34^. Interestingly, *Meox1* transcripts persisted at a high level at E10.5 in cluster 0, where *Hoxa2* was up-regulated in STZ embryos. Furthermore, *Postn*, a direct target gene of *Meox1*^35^ normally expressed in PA1 to dictate tooth development^36^, was also upregulated in these cells (**Extended Data Fig. 4c**). Other than dysregulation of genes related to PA patterning and development, the top GO terms enriched in cluster 0 DEGs in STZ were associated with cell migration, and these results are consistent with DAR findings. (**Fig. 2l, Extended Data Fig. 4e, f**). Taken together, multi-modal analysis of neural crest cells at single cell resolution suggests that a subset of PA2 neural crest cells that are relatively undifferentiated are modified epigenetically by PGDM, resulting in transcriptional dysregulation of genes related to patterning, cell migration and cellular differentiation.

## A novel *Alx3* positive AHF subpopulation specifically affected by PGDM

We performed a similar characterization of mesodermally-derived cardio-pharyngeal progenitor cells and cardiomyocytes, as they also displayed a large number of DARs upon hyperglycemia. Upon re-clustering this population using scRNA-seq, we detected five cardiomyocyte sub-types and ten mesodermal progenitor sub-types (**Fig. 3a, b** and **Extended Data Fig. 5a**). The integration of scRNA-seq with scATAC-seq data (**Fig. 3c** and **Extended Data Fig. 5b**) revealed that the C15 cluster in scATAC-seq, which displayed the largest number of DARs, corresponded to one of two anterior heart field cell (AHF) clusters, which we labeled AHF2. AHF1 and AHF2 had very similar transcriptional profiles, but AHF2 displayed lower expression of AHF marker gene *Hand2* and near absence of *Rgs5* and *Armh4* (**Fig. 3b, d and Extended Data Fig. 5c**).

**Fig. 3.**
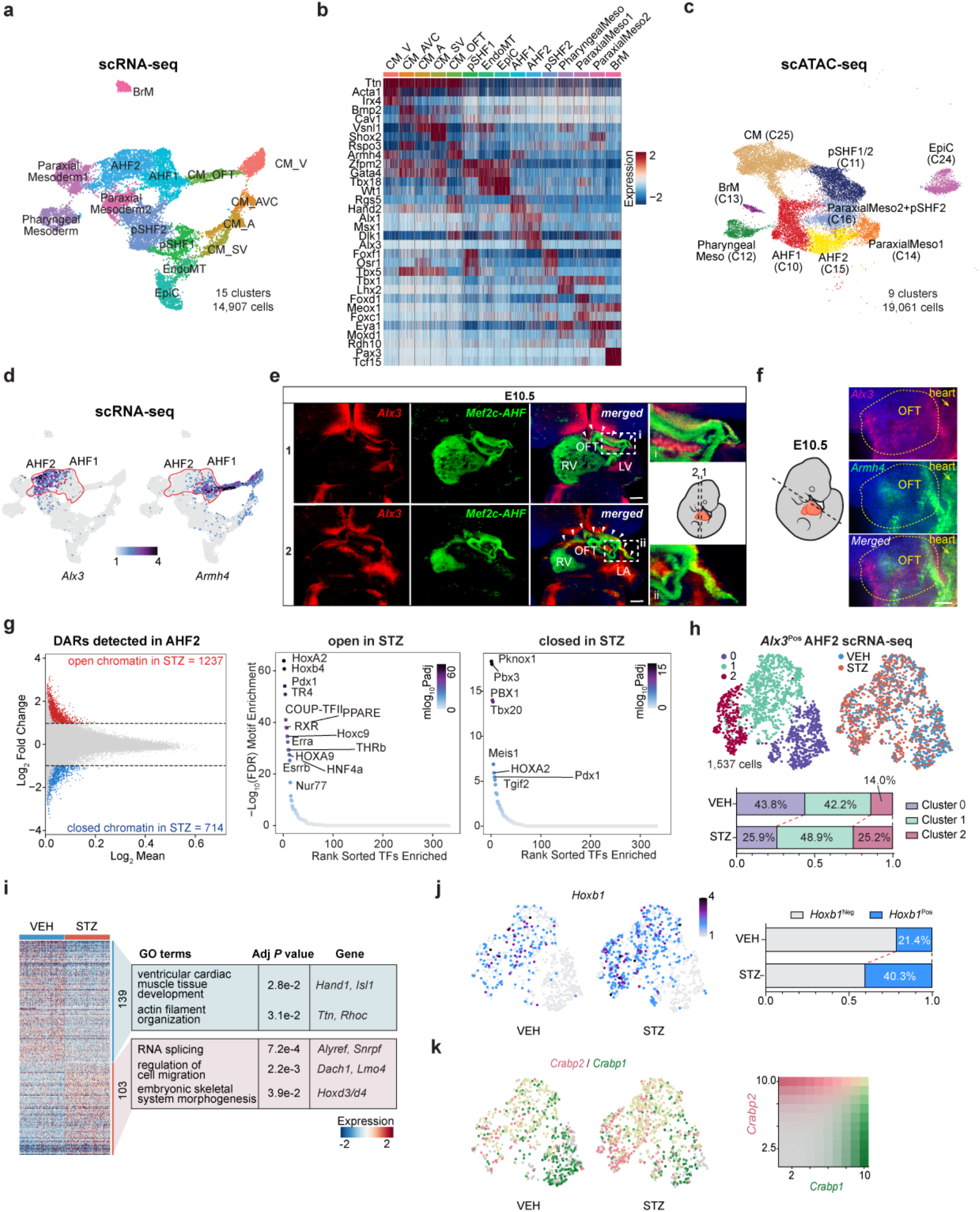
Maternal Diabetes Epigenetically Alters *Alx3*-Positive Anterior Heart Field Cells with Disruption of Anterior-Posterior Patterning. **a**, scRNA-seq UMAP representation of mesodermal cell subset population (Meso/CPP”, “Cardiomyocyte”, or “Epicardium” in Figure 1b), including both VEH (n=3) and STZ (n=3), colored by cluster identity. **b**, Heatmap of expression levels of top marker genes for each sub-cell-type in the mesodermal cell population. Scale indicates z-scored expression values. All genes represented have an adjusted p-value smaller than 1e-10 (Wilcoxon rank-sum test). **c**, scATAC-seq UMAP representation of mesodermal cell population, including both VEH (n=3) and STZ cells (n=3), colored by cluster identity. **d**, Expression pattern of *Alx3* (left) and *Armh4 (right)*. AHF1 and 2 are circled in red. Scale bar indicates z-scored expression values. **e**, Optical sections from the same data as Extended Data Movie 1, with angle of section indicated in cartoon (section 1, top panels from a more ventral slice; section 2, bottom panels from a more dorsal slice). *Alx3* (red), Mef2c-AHF (green) and DAPI (blue) are shown. The scale bar represents 100 µm. **f**, Optical sections of outflow tract region (circled by yellow dotted line) from the same data as Extended Data Movie 2, with angle of section indicated in cartoon. *Armh4* (green, top), *Alx3* (red, middle), and DAPI (blue) are shown. The scale bar represents 50 µm. OFT, outflow tract. **g**, MA plot of differentially accessible regions (DARs) between VEH and STZ (left) in AHF2 population. Red and blue dots represent more or less accessible chromatin regions, respectively in STZ (FDR <= 0.05 & Log2FC >= 1). Enriched TF binding motifs in more accessible open (middle) or less accessible closed (right) DARs in AHF2 population from STZ embryos. **h**, scRNA-seq UMAP representation of *Alx3* expressing (*Alx3*^Pos^) AHF2 cells colored by cluster identity (upper left) or by conditions (upper right). Population distribution normalized to total number of cells per each sample in *Alx3*^Pos^ AHF2 cells. Each number inside the bar plot (bottom) represents the percentage of the corresponding cell type among the total cell number of each condition. Statistics of STZ vs VEH performed by permutation test in scRNA-seq data for cluster 0, FDR<0.0005, Log2FD=-0.74; for cluster 1, FDR<0.01, Log2FD=0.22; for cluster 2, FDR<0.0005, Log2FD=0.78. **i**, Heatmap of differentially expressed genes (DEGs) between VEH or STZ *Alx3*^Pos^ AHF2 cells. All the detected DEGs with adjusted p-val < 0.05 and Log2FC > 0.25 are shown. Top GO terms enriched in upregulated or downregulated DEGs are shown with representative genes. Scale bar indicates z-scored expression values. **j**, Expression of *Hoxb1* on UMAP of *Alx3*^Pos^ AHF2 cells. Scale bar indicates z-scored expression values. Percentage of *Hoxb1* positive cells to the total cell number in each condition (right). Statistics of STZ vs VEH performed by permutation test in scRNA-seq data for *Hoxb1* positive cells, p=0.0001, Log2FD=0.82. **k**, Expression of *Crabp1* (green) or *Crabp2* (red) on UMAP of *Alx3*^Pos^ AHF2 cells. Scale grid indicates z-scored expression values for each gene. CM_V, ventricular cardiomyocyte; CM_AVC, atrioventricular canal cardiomyocyte; CM_A, atrial cardiomyocyte; CM_SV, sinus venosus cardiomyocyte; CM_OFT, outflow tract cardiomyocyte; pSHF1/2, posterior second heart field 1/2; EndoMT, endothelial mesenchymal transition; EpiC, Epicardium; AHF1/2, anterior heart field 1/2; PharyngealMeso, pharyngeal mesoderm; ParaxialMeso1/2, paraxial mesoderm 1/2; BrM, branchiomeric muscle

This AHF2 population was clearly distinguishable from pSHF cells, which are marked by *Foxf1* and *Osr1* (**Fig. 3a, b**). The transcription factor *Alx3* was specifically expressed in the AHF2 population, uniquely marking this newly recognized subset of the AHF (**Fig. 3d**). To determine if the AHF2 cells are truly of AHF origin, we examined overlap of *Alx3* expression with AHF-derived cells marked by Cre-recombinase under control of the Mef2c AHF-specific enhancer (Mef2c-AHF-Cre: Ai6 AHF lineage tracing mouse model)^37^. RNA *in situ* hybridization with light sheet microscopy revealed that *Alx3*-positive cells were distributed in a continuum from the distal outflow tract towards the posterior bilateral sides of the embryo. Of these, the more anterior *Alx3*-positive cells overlapped with the Mef2c-AHF-Cre lineage, demonstrating that *Alx3* marks a subset of the AHF cells (**Fig. 3e and Extended Data Movie 1**). We further found that *Alx3*-expressing cells resided posterior to the *Armh4*-expressing AHF1 cells, abutting the AHF1 cells of the distal outflow tract (**Extended Data Fig. 6a, b and Extended Data Movie 2)**. Optical section demonstrated that the *Alx3*-expressing cells and the *Armh4*-expressing cells share similar AHF regions at the distal outflow tract region without overlapping one another **(Fig. 3f)**. Thus, PGDM affects a very specific subset of the SHF that appears to lie between the classically recognized AHF and pSHF.

To investigate the *Alx3*-enriched subpopulation of the SHF, we analyzed the mouse cardiac development single cell atlas data from E7.75 to E9.25 previously generated by our lab^38^. *Alx3*-positive cells (*Alx3*^Pos^) were detectable within clusters annotated as AHF, with significant expression detected in a small number of cells by E9.25 (**Extended Data Fig. 6c-e**). DEG analysis between *Alx3*^Pos^ and *Alx3*^Neg^ AHF cells at E9.25 demonstrated that *Alx3*^Pos^ AHF cells expressed higher levels of genes related to epithelial-to-mesenchymal transition and skeletal system morphogenesis, with lower levels of genes involved in cardiomyocyte development (**Extended Data Fig. 6f**). By E10.5, the *Alx3*^Pos^ AHF cells expanded and although the number of DEGs between AHF2 and AHF1 increased considerably, they still displayed similar GO terms (**Extended Data Fig. 6g, h)**.

DARs within AHF2 cells with increased accessibility in STZ compared to VEH were enriched in GO terms related to anterior/posterior specification, mesenchymal development, and mesonephros development. In contrast, DARs with reduced accessibility were enriched in terms related to Wnt signaling, maintenance of cell number, differentiation of cardiac muscle, and heart development (**Extended Data Fig. 7a**). TF motif analysis in DARs with increased accessibility in PGDM/hyperglycemia revealed enrichment for HOX gene groups (Hoxa2, Hoxb4, Hoxc9, Hoxa9), as well as COUPTFII and RXR, which are involved in establishing pSHF cells, and are not normally accessible in AHF cells^39,40^. In contrast, motifs enriched in DARs with decreased accessibility in PGDM/hyperglycemia included PBX1, MEIS1 and PKNOX1, which normally function together to activate more anterior Hox genes^40,41^; and disruption of *Pbx* genes has been reported to cause outflow tract malformation both in mice and human^42^. In addition, motifs for TBX and TGIF, both involved in cardiac differentiation^43^, were enriched in closed DARs (**Fig. 3g**). Thus, the AHF2 cells, which at baseline are more posterior to AHF1 cells and have a decreased cardiomyocyte signature, appeared to be even more “posteriorized” and less cardiac-like in offspring of STZ-treated mothers.

To investigate the effects of PGDM/hyperglycemia more precisely on *Alx3*^Pos^ AHF2 cells, we re-clustered the *Alx3*-expressing AHF2 cells and detected three distinct sub-clusters (**Fig. 3h**). Marker genes for Cluster 0 were enriched for genes related to cardiomyocyte development, including cardiac TFs like *Hand1/2, Isl1*, and *Tbx2/5*, while those for Cluster 1 and 2 were enriched for genes involved in cell migration, receptor protein tyrosine kinase signaling, and skeletal system morphogenesis (**Extended Data Fig. 7b**). Cluster 1 and 2 cells were increased in number under hyperglycemia, while Cluster 0 numbers were decreased compared to control (**Fig. 3h**). These findings suggest that PGDM/hyperglycemic conditions led to a cellular state transition from Cluster 0 towards Cluster 1 and 2, characterized by further loss of cardiac TF expression and gain of a skeletal-like state. This was further confirmed by DEG analysis between the STZ and VEH groups in *Alx3*^Pos^ AHF2 cells (**Fig. 3i**). Notably, the pSHF Hox gene, *Hoxb1*,^40^ was expressed almost exclusively within clusters 1 and 2 where STZ cells increased, and was nearly absent in cluster 0, whereas STZ treatment led to both significant increases in the number of *Hoxb1* positive cells and its expression levels (**Fig. 3j**). This was corroborated by the STZ-mediated increase of the chromatin accessibility of the distal regulatory element of *Hoxb1*, calculated as a “peak-to-gene link” by ArchR. (**Extended Data Fig. 7c**).

Cluster 2 was notable not only for *Hoxb1* expression among the STZ group, but also for decreased expression of *Crabp1* and increased *Crabp2* transcripts, competing modulators of retinoic acid signaling (**Fig. 3k**). *Crabp1*, normally highly expressed in the AHF, enhances retinoic acid catabolism, while *Crabp2*, which is expressed at higher levels in the pSHF, facilitates nuclear transport of retinoic acid and subsequent transcriptional activation, resulting in posteriorization of SHF progenitors^44^. This raised the possibility that, under PGDM/hyperglycemia, abnormal anterior-posterior patterning may occur in the *Alx3*^Pos^ AHF2 due to ectopic retinoic acid-induced posteriorization, including induction of *Hoxb1* expression.

## Retinoic acid signaling is dysregulated in cell types affected by PGDM

Given the dysregulated *Crabp1* and *Crabp2* expression and anterior-posterior patterning defects observed in the STZ group, we specifically evaluated RA signaling in cells from VEH or STZ treatment. Using ChromVAR analysis, which can assess enrichment of specific TF activity at the single cell level, we found that PGDM drove significant increased enrichment for RA signaling-related TFs (RAR and RXR) in most cell types, including PA2 neural crest-derived cells and AHF2 cells (**Extended Data Fig. 8a, b and Extended Data Table 2**). This was in agreement with the TF motif enrichment analysis in DARs, which showed that the RXR motif and its downstream effectors’ motifs, such as COUP-TFII and HOX, were highly enriched in more accessible DARs under PGDM/hyperglycemia in PA2 and PA4/6 neural crest and AHF2 cells (**Fig. 2h, 3g, and Extended Data Fig. 3h**).

To investigate mechanisms by which altered RA signaling may contribute to transcriptional and epigenomic changes in hyperglycemia, we selected distal regulatory regions with differential accessibility that had peak-to-gene links with differentially expressed genes between VEH and STZ, and also had RA signaling-related motifs within the scATAC-seq peaks. Among these, we focused on several candidate enhancers for dysregulated genes in PA2 neural crest cells, with further experimentation related to *Tfap2a* and *Nr2f1* gene activation (**Fig. 4a**). *Tfap2a* is implicated in neural crest cell induction and migration, and together with *Nr2f1*, binds to active enhancers that are critical for maintaining cranial neural crest cells in an undifferentiated state^30^. We performed luciferase reporter assays using a validated RA-responsive *Cyp26a1* enhancer element as a positive control and found that high glucose itself did not enhance transcription, suggesting that heightened RA signaling is not a direct consequence of hyperglycemia (**Fig. 4b**). We further found PA2 neural crest candidate regulatory elements for *Tfap2a* and *Nr2f1* were able to activate transcription in the presence of RAR/RXR receptors and showed further modulation by addition of RA. RA-dependent activation was abolished upon deletion of putative RAR/RXR binding sites (**Fig. 4c, d**). The dysregulation of RA signaling present in STZ embryos was supported by analysis of the data using Weighted Correlation Network Analysis (WGCNA)^45^. This approach enables identification of co-variate transcriptional modules, gene regulatory networks (GRN), across individual cells without the influence of possible bias introduced through sub-clustering processes and manual annotation of the neural crest compartments. WGCNA revealed a PGDM-induced abnormality in a GRN that was induced only in PA2 of STZ embryos. In particular, *Tfap2a* downstream genes such as *Eya1, Dlx5/6*, and *Hoxa2* constituted a core network within this GRN, consistent with PA2-specific aberrant RA signaling leading to increased *Tfap2a*/*Nr2f1* and subsequent patterning defects in the setting of PGDM (**Extended Data Fig. 8c, d**).

**Fig. 4.**
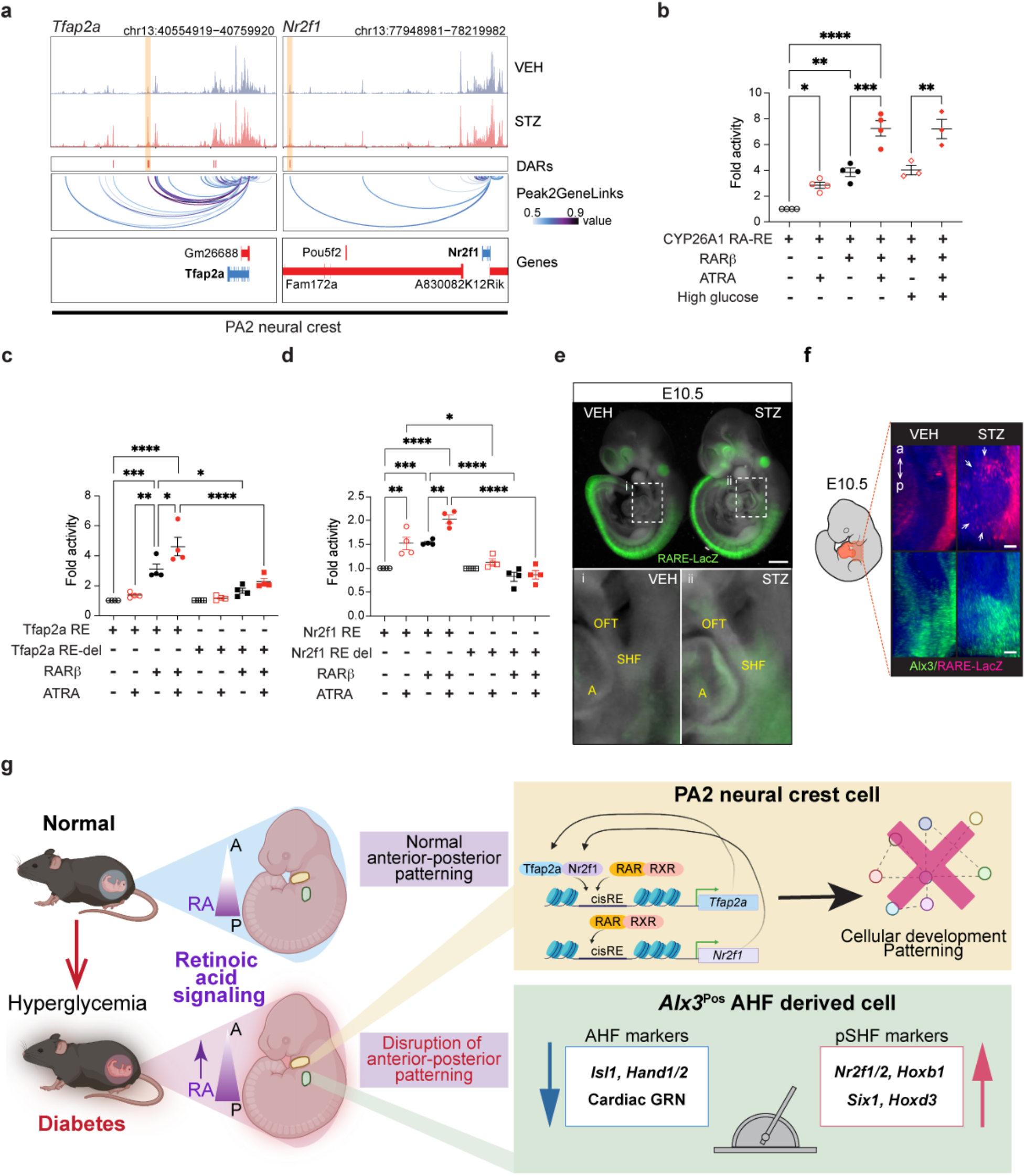
Disruption of Retinoic Acid Signaling Associated with Pharyngeal Arch and *Alx3*^pos^ Anterior Heart Field Anterior-Posterior Patterning Defect. **a**, Genome browser plots for the *Tfap2a* or *Nr2f1* locus. The top two rows represent the chromatin accessibility in VEH or STZ. The third track shows genomic locations of detected DARs with more accessibility in STZ (red rectangles). The DARs selected for further analyses are highlighted in yellow boxes. The second track from the bottom represent the links between peaks and gene (“Peak2GeneLinks”), calculated by ArchR. Darker lines represent stronger links. The bottom track shows the gene location and transcriptional direction (red – positive strand; blue – negative strand). **b**, Relative luciferase activity driven by the known RA response elements (RA-RE) for hCYP26A1 with exogenous RAR*β* transduction, addition of 1µM ATRA and high glucose condition (25 mM). **c**, Relative luciferase activity driven by the candidate regulatory region for *Tfap2a* with (RE) /without deletion (RE-del) of the putative RAR/RXR binding site under conditions indicated. **d**, Relative luciferase activity driven by the candidate regulatory region for *Nr2f1* with (RE) /without deletion of the putative RAR/RXR binding site under conditions indicated. **e**, Whole mount RNA *in situ* hybridization for LacZ transgene driven *in vivo* by a retinoic acid response element (RARE-LacZ) in E10.5 VEH or STZ embryos. Regions highlighted by the white dashed squares are shown in (i) and (ii). Scale bar represents 500 µm. OFT, outflow tract; SHF, second heart field; A, atrium. **f**, Representative images of whole mount RNA *in situ* hybridization of the SHF area of E10.5 embryos in lateral view using light sheet microscopy. *RARE*-*LacZ* (magenta, top) and *Alx3* (green, bottom) are shown for VEH or STZ embryos. Still images were extracted from Extended Data Movie 3. White arrows indicate ectopic LacZ signals in *Alx3* positive area. The scale bars represent 50 µm. The cartoon illustrates the approximate location shown in the images. **g**, Model of PA2 neural crest and *Alx3*^pos^ AHF-specific epigenomic and transcriptional alterations and their consequences through aberrant RA signaling in maternal diabetes. *p<0.05, **p<0.005, ***p<0.001, and ****p<0.0001 by Tukey’s multiple comparisons test after ANOVA.

Similarly, WGCNA using the mesodermal cardio-pharyngeal cells identified a GRN with significant module activity score changes between VEH and STZ samples only in the AHF2 population (**Extended Data Fig. 8e, f**). This module contained genes related to cardiac outflow tract and right ventricle development, including critical regulators such as *Hand1/2, Isl1, Bmp4*, and *Gata6*, which were downregulated in STZ as members of a core network of the GRN. This is consistent with the diminished “anterior” nature of the *Alx3*^Pos^ AHF2 in offspring of PGDM mothers. To experimentally determine if RA signaling activity, normally present in the pSHF, was ectopically enhanced in AHF2 cells within STZ-treated embryos, we exposed a cohort of transgenic female mice containing a LacZ reporter driven by a RA response element (RARE-LacZ) to STZ to establish a PGDM model in this genetic background. As expected, direct detection of LacZ expression in VEH offspring using RNAscope indicated RA activity was evident in the pSHF region, but not in Alx3-expressing cells. In contrast, STZ-treated offspring demonstrated an enlarged LacZ-positive domain with marked anterior extension compared to the VEH control samples (**Fig. 4e, f and Extended Data Movie 3**). This result provided *in vivo* evidence that “posteriorization” of the *Alx3*^Pos^ AHF2 population occurs with anterior expansion of RA signaling in offspring of mothers with PGDM, likely contributing to impaired outflow tract development.

## Discussion

In this study, we used single cell multi-modal analyses to uncover mechanisms by which environmental influences such as PGDM can cause developmental anomalies. We demonstrated selective epigenomic vulnerability of a small subset of cells that help explain cardiac and craniofacial defects that are observed in PGDM despite all embryonic cells experiencing similar environmental exposure. Specifically, we found that a previously unrecognized subpopulation of the *Alx3*^Pos^ AHF is uniquely “posteriorized” in the setting of PGDM. Furthermore, a subset of relatively less differentiated neural crest cells within PA2 and PA4/6 also had evidence of abnormal cell differentiation and anterior-posterior patterning. Analyses of regulatory elements in these specific cell types demonstrated that retinoic acid signaling and downstream *Hox* genes were dysregulated in AHF2 and PA2, with ectopic retinoic acid activity resulting in disruption of proper patterning and loss of cell type specific transcription signatures (**Fig. 4g**). Mechanistically, it appears RA metabolism dysregulation by altered *Crabp1* and *Crabp2* expression contributes to the alterations in AHF2, while RXR-mediated transcriptional upregulation of *Tfap2a* and *Nr2f1* disrupts PA2 differentiation and patterning. In particular, the presence of putative binding sites of *Tfap2* and *Nr2f1* in the Rxr-dependent *Tfap2a* regulatory element suggests a positive feedback mechanism reinforcing RA signaling-mediated transcriptional dysregulation under PGDM/hyperglycemia in PA2 neural crest cells (**Fig. 4g**).

These findings begin to address how an environmental influence can affect specific embryonic cell types and lead to unique developmental phenotypes and congenital birth defects. In the future, it will be important to understand why certain cell types are more vulnerable than others and how environmental influences as occur in PGDM cause the very specific epigenomic shifts described here. While PGDM clearly causes shifts in RA signaling, the precise mechanism by which regulators of RA metabolism and RXR-dependent transcription are altered will require further study. For example, increased levels of beta-hydroxybutyrate present in ketotic mothers with PGDM may contribute to specific epigenomic alterations through histone modification in the embryo^13,14^; future advances that enable interrogation of individual epigenomic marks in single cells will help explore this possibility. Ultimately, discovery of the consequences of environmental factors on embryonic development using multi-modal single cell genomics will not only lead to potential gene-environment combinations that may contribute to the complex and multifactorial diseases, but also preventive approaches to protect vulnerable cells.

## Methods

### Animal

Animal studies were conducted in compliance with all relevant ethical regulations in the animal use protocols, UCSF animal use guidelines and the NIH Guide for the Care and Use of Laboratory Animals. All protocols relating to animal use were approved by the Institutional Animal Care and Use Committee (IACUC; AN189140) at UCSF which is accredited by the Association for Assessment and Accreditation of Laboratory Animal Care (AAALAC). C57BL/6J wild type (JAX stock no. 000664), RARE-hsp68LacZ (JAX stock no. 008477), and B6.Cg-Gt(ROSA)26Sortm6(CAG-ZsGreen1)Hze/J (Ai6) (JAX stock no. 007906) mice were purchased from Jackson Laboratory (Bar Harbor, ME), Mef2c-AHF-Cre mice^1^ were provided by Brian Black’s lab. All animals were housed in a 12-hour light/dark cycle and given ad libitum access to standard chow and water unless otherwise documented.

### Model of maternal diabetes mellitus

Diabetes induction was performed using streptozotocin (STZ) (Sigma-Aldrich, S0130) as previously described with minor modifications^2^. Diabetes mellitus was induced in ∼10-week-old female C57BL/6 mice or RARE-hsp68LacZ mice by intraperitoneal injections of 60 mg/kg body weight STZ for 5 consecutive days. Two weeks after the last STZ injection, blood glucose levels were measured after 6 hr fasting using a Zoetis, AlphaTRAK2 glucometer. Mice were diagnosed as diabetic if blood glucose measurements were greater than or equal to 250 mg/dL and were subsequently bred to 12-week-old C57BL/6 male mice. Pregnant females were identified by echocardiography performed at E6.5 or E7.5 and euthanized to harvest embryos at E10.5 for single-cell RNA sequencing, single-cell ATAC sequencing, and whole-mount in situ hybridization experiments, and at E18.5 for histological examination.

### Embryo dissection, single-cell library preparation and sequencing

C57Bl6/J E10.5 mouse embryos were dissected in cold PBS (Life Technologies, 14190250) with 1%FBS (ThermoFisher Scientific, 10439016) and truncal region was further microdissected to include the heart tube region, the SHF and mesodermal regions behind the heart tube, as well as the second pharyngeal arches (**Extended Data Fig. 2a**). Three embryos with matched somite counts each were collected from females in the VEH or STZ treatments. Dissected cardiac tissue was incubated in 200 µl TrypLE (ThermoFisher Scientific, 12563029) for 5 min, triturated with a 200-µl pipette tip, and incubated for an additional 5 min. The TrypLE solution was quenched with 600 µl PBS with 1% FBS. Cells were filtered through a 70-µm cell strainer (BD Falcon, 08-771-2), centrifuged at 150g for 3 min, and resuspended in PBS with 1% FBS. The cell suspension was prepared by adjusting the cell number to recover 10K cells per sample according to the manufacturer’s instructions (Chromium Single Cell 3’ Reagent Kit v.3, 10X Genomics). GEM generation for scRNA-seq was performed on the Chromium controller. The remaining cells were then subjected to nuclear isolation according to the 10x nuclei isolation protocol (Nuclei Isolation for Single Cell ATAC Sequencing, CG000169, 10X Genomics). The nuclei solution was adjusted so that the target nuclei number was 10K per sample (Chromium Next GEM Single Cell ATAC Reagent Kits v1.1, 10X Genomics), followed by transposition reaction. GEM generation for scATAC-seq was performed using the Chromium controller (10X Genomics).

The subsequent scRNA-seq and scATAC-seq library preparations were performed according to the protocols described above using Chromium Single Cell 3’ Reagent Kits v3 (PN-1000075) and Chromium Next GEM Single Cell ATAC Reagent Kits v1.1 (PN-1000176), respectively. Additionally, Chromium Chip B Single Cell Kit (PN-1000153) and Chromium i7 Multiplex Kit (PN-120262) were used for scRNA-seq library preparation, and Chromium Next GEM Chip H Single Cell Kit v1.1 (PN-1000161) and Single Index Kit N Set A (PN-1000212) were used for scATAC-seq library preparation. The scRNA-seq libraries were sequenced using HiSeq4000 (Illumina) and NovaSeq (Illumina), and the scATAC-seq libraries were sequenced using NovaSeq, in accordance with the manufacturer’s protocol.

### scRNA-seq analysis

Raw sequencing data were preprocessed with the Cell Ranger v.5.0.0 pipeline (10X Genomics). Data from all samples was merged by cellranger-aggr in the pipeline described above and normalized to the same sequencing depth, resulting in a single gene-barcode matrix. Further analysis was performed using Seurat v4.0 R package^3^ with reference to the Seurat web tutorials. Low quality cells were removed from analyses, keeping the cells with number of genes per cells between 4,000 and 9,500, with a UMI count per cell between 10,000 and 100,000, with mitochondrial gene percentage between 1 and 6%, and with ribosomal gene percentage between 10 and 30%. After the filtering step, for normalization, the data were transformed using SCTransform function, setting mitochondria gene percentages and ribosomal gene percentages as a variance to be regressed. Then, principal component analysis (PCA) was performed with RunPCA. Batch correction among each embryo was performed using Harmony^4^. Cells were clustered using top 32 principal components and visualized using a Uniform Manifold Approximation and Projection (UMAP) dimensionality reduction (RunUMAP, FindNeighbors, and FindClusters). For clustering, a vector of resolution parameters was passed to the FindClusters function and the optimal resolution that established discernible clusters with distinct marker gene expression was selected. To identify marker genes, the clusters were compared pairwise for differential gene expression using the FindAllMarkers function (min.pct = 0.45, logfc.threshold=0.4, and return.thresh (p-value cut-off) = 1 × 10^−10^). By cross-referencing the resulting cluster-specific marker genes with known cell type-specific maker genes, we classified them into clusters of seven cell types. The numbers of cells in each of those major cell type clusters in each condition (VEH vs STZ) were then calculated and plotted in a bar plot using the ratio of each cluster to the whole cells in each condition. Cardiomyocyte and Meso-CardiopharyngealProgenitors clusters, NeuralCrest cluster, PA2 neural crest cluster, and AHF2 cluster were re-clustered for the further analyses. Clustering was undertaken as previously described with minor modification. For the SCTransform step, in both subset analyses, we used mitochondria gene percentage and cell cycle scores as a variance to be regressed. In the mesoderm-cardiomyocyte subset analyses, top 30 PCAs were used for dimensional reduction. In the neural crest cell subset analyses, top 34 PCAs were used for dimensional reduction. In the PA2 neural crest cell subset analyses, top 22 PCAs were used for dimensional reduction. In the AHF2 cell subset analyses, top 30 PCAs were used for dimensional reduction. Differential gene expression analysis between conditions (VEH and STZ) in each sub cell types was performed using the FindAllMarkers function. For visualization of gene expression data, the following Seurat functions were used: FeaturePlot, VlnPlot, DotPlot, and DoHeatmap. Enrichr^5^ was used for gene ontology (GO) enrichment analysis. All the indicated p-values for the GO term analyses are adjusted p-values calculated with Fisher exact test.

### Gene regulatory network analysis

WGCNA (v1.70-3) ^6^ was used to identify gene regulatory networks which were dysregulated under hyperglycemic conditions. Transcriptional data for either subset of scRNA seq data subject to analysis were retrieved via Seurat’s GetAssayData function, returning a cell by gene expression matrix. This matrix was processed with WGCNA’s blockwiseModules function (minModuleSize = 10, mergeCutHeight = 0.15), and the resultant modules’ gene activity in each cluster was subsequently scored with the AddModuleScore function in the source Seurat object. Statistical significance of the difference in the mean module scores between treatment conditions within cells of a given cluster of interest was assessed using a linear mixed-effects model (lmerTest v3.1-3) with Benjamini-Hochberg multiple-testing correction applied to resultant p-values (stats v4.0.5). The mouse id from which each cell is derived is modeled as a random effect. Statistically significant modules were subsequently input to the StringDB web-interface (www.string-db.org) to compute, analyze, and visualize protein-protein interaction networks.

### scATAC-seq analysis

Raw sequencing data were preprocessed with the Cell Ranger ATAC v.2.0 pipeline (10X Genomics). Subsequent analyses were performed using ArchR v1.0.1 R package^7^ with reference to the ArchR web tutorials. Low quality cells were removed based on TSS enrichment score and the number of fragments per nuclei (createArrowFiles function with minTSS=7, minFrags=19952, maxFrags=1E+6), and nuclei doublets were removed through addDoubletScores and filterDoublets functions, resulting in making an ArchRproject to be analyzed. Through the genome-wide tiling with iterative Latent Semantic Indexing, the dimensionality reduction was performed using the addIterativeLSI function. Batch correction among each embryo was performed using the addHarmony function. Then, clustering and the 2D embedded visualization in UMAP space were performed using the addClusters function, and the addUMAP and plotEmbedding functions, respectively. Gene scores (GS), which were calculated by the accessibility of promoter and gene body regions of each gene and can be treated as a proxy of expression levels of a corresponding gene, were extracted to identify the cluster features using the getMarkerFeatures with useMatrix=“GeneScoreMatrix”. For peak calling per cluster, the addGroupCoverages and the addReproduciblePeakSet functions with peakMethod = “Macs2” were used. Furthermore, to identify the cluster specific feature peaks, the getMarkerFeature functions with useMatrix=“PeakMatrix” was used, and transcription factor (TF) binding site motif enrichment analysis on the resultant peaks was performed using HOMER^8^. For differential accessible region (DAR) analysis, peak calling was performed for each condition (VEH or STZ) in each cluster and differential analyses were performed using the getMarkerFeatures function by setting a cluster from STZ as useGroups and a corresponding cluster from VEH as bgdGroups in the function. The statistically significant DARs were defined with FDR less than 0.05 and Log2 fold change greater than 1. Motif enrichment analyses on the detected DARs were performed using the peakAnnoEnrichment function. To integrate the scRNA-seq data from the same fetus heart samples, the aforementioned Seurat object with whole cell types were preprocessed with the FindVariableFeatures from the Seurat package to extract variable genes and transformed by the as.SingleCellExperiment function from Seurat packages. Then, the Seurat object was integrated using the addGeneIntegrationMatrix with useImputation=F. Considering the marker GS, marker peaks and the results of scRNA-seq data integration, the found clusters were annotated. Integration with scRNA-seq data was validated by calculating prediction scores, integration scores during integration, or Jaccard indices between the scATAC-seq annotation and the scRNA-seq annotation^9^. To assess enrichment of TF activity per cell, ChromVAR analyses for retinoic acid signaling related receptor TFs were performed using the addDeviationsMatrix and getVarDeviations functions. The differentialDeviations function was used to determine whether there is a significant difference between the bias-corrected deviations for a TF motif between conditions per each cell type ^10^. To identify an association of each peak and expression of a corresponding gene per cluster, the addPeak2GeneLinks function was used. The found associations between peak to gene were integrated with the differential expressed genes in the same cluster between conditions VEH and STZ to identify the candidate enhancers for those DEGs.

### Single cell proportion test

For statistical tests addressing cell type population changes between conditions, a permutation test is used to calculate a p-value for each cluster implemented in the R package scProportionTest with “n_permutations = 10,000” (https://github.com/rpolicastro/scProportionTest/releases/tag/v1.0.0). scRNA populations were analyzed directly from Seurat as described in package documentation, and scATAC populations were analyzed by manually exporting ArchR metadata to scProportionTest.

### Peak annotation

ChIPpeakAnno R library^11^ was used for converting a bed file of detected peaks out from ArchR pipeline to a GRanges format. Then, the annotatePeak function from ChIPseeker R library^12^ was used for annotating genome location with “tssResion” augment setting as 2 kb. The plotDistToTSS function was used for visualizing the location of each peak from TSS.

### GREAT analyses

Inferring the functional significance of cis-regulatory regions found in scATAC-seq analyses was performed with GREAT v4.0.4 webtool^13^ (http://great.stanford.edu/public/html/index.php) using default parameters. Detected regions were filtered with a significant threshold set at an adjusted p-value less than 0.05 and Log2 fold change greater than 1.

### *in situ* hybridization experiments

Whole mount RNA in situ hybridization was performed in E10.5 embryos using Multiplex Fluorescent Reagent Kit v2 (Advanced Cell Diagnostics, 323100) as previously reported^14^. Each experiment was repeated using at least three biological replicates to determine the spatial expression of the genes. The *in situ* hybridization probes used in this study are as follows: Mm-Alx3-C2 (Advanced Cell Diagnostics, 1127551-C2), ZsGreen-C3 (Advanced Cell Diagnostics, 461251-C3), Mm-Armh4-C1 (Advanced Cell Diagnostics, 1085041-C1), and Ecoli-LacZ-C3 (Advanced Cell Diagnostics, 313451-C3). Whole-mount embryos were imaged in 0.1% PBST using the Leica MZ FLIII fluorescence stereomicroscope (acquisition software LAS v.3.7.4).

For light sheet microscopy imaging, the whole mount samples were microdissected to keep the area of interest and were mounted in 1.5% low melting temperature agarose (Fisher Scientific, Hampton, NH) in a glass capillary with matching piston rod (Sigma Aldrich, Steinheim). Samples were then cleared in EasyIndex optical clearing solution with a refraction index of 1.465 (Lifecanvas technologies, Cambridge, MA) at 4 °C for two nights minimum before imaging, by extending the agarose-embedded sample into the solution. Samples were then imaged on a Zeiss Z.1 Lightsheet microscope. Images were processed with Fiji and Imaris.

### Luciferase assay

Candidate enhancer sequences identified from the integrated analysis from scRNA-seq and scATAC-seq were PCR amplified (TAKARA, PrimeSTAR GXL) from mouse genomic DNA and PCR products cloned into pGL4.23 (Promega) pre-digested with XhoI and HindIII using Cold Fusion Cloning kit (System Biosciences). Deletion of putative TF binding sites from those enhancers were performed using QuickChange II XL site-directed mutagenesis kit (Agilent Technologies). All constructs were confirmed by Sanger Sequencing (Quintara Biosciences). The primers used for cloning enhancer sequences and establishing deletion mutants are listed in the Extended Data Table 3. For the luciferase assay, 7×10^4^ HeLa cells were seeded per-well in 24 well plates 24 hours before transfection. All enhancer or mutant enhancer reporter constructs (200ng) were transfected with 20ng of pRL (Renilla luciferase control vector, Promega) and either empty vector (pcDNA3.1, Thermofisher Scientific) or hRARß-RXRa (Addgene, #135415) ^15^ with 2.4ul Fugene HD (Promega) in opti-MEM (Gibco). All transfectants were stimulated with either vehicle (DMSO) or 1 μM ATRA for 8 hours starting 24 hours after transfection. Each construct was transfected in quadruplicate and repeated at least three times. Luciferase assays were conducted 8 hours after stimulation using a dual luciferase kit (Dual-Luciferase® Reporter Assay System, Promega) according to manufacturer’s protocol.

Luciferase activity was measured in SpectraMax MiniMax 300 imaging cytometer with SoftMax Pro6, Version 6.4 (Molecular Devices). pGL3-Basic-hCYP26A1-FL-luciferase (Addgene, #135566) ^16^ was used as a positive control in the assay system. The relative luciferase activity of each construct (arbitrary unit) was reported as the fold changed compared with empty vector.

### Statistics

Statistical analyses were performed using GraphPad Prism 9. The number of replicates, statistical test used, test result, and the significance levels are described in each figure legend. For all the quantification, the mean ± s.e.m. is presented. No experimental sample was excluded from statistical analysis. When presenting representative results showing a gene expression pattern in a wild-type embryo, at least three independent embryos were analyzed.

## Data availability

All the sequencing data have been deposited in NCBI’s Gene Expression Omnibus and are accessible through GEO series accession number GSE198905.

## Code availability

All analyses were performed using standard protocols with previously described R packages. All R scripts are available on GitHub (https://github.com/SrivastavaLab-Gladstone/Nishino_DM_2022).

## Acknowledgements

We thank members of the Srivastava laboratory for discussion and feedback; Bethany Taylor for editorial and graphics assistance; Giovanni Maki for graphics assistance; and Kathryn Claiborn for editorial review. We acknowledge the center for advanced technology (CAT) for sequencing; the Gladstone Histology and Light Microscopy Core for their technical support and the Gladstone Animal Facility for support with mouse colonies. D.S. is supported by National Institutes of Health/NHLBI (P01 HL146366, R01 HL057181, R01 HL015100, R01 HL127240), Roddenberry Foundation, L.K. Whittier Foundation, Dario and Irina Sattui, Younger Family Fund, and Additional Ventures. T.N. is supported by the Japan Society for the Promotion of Science Overseas Research Fellowship. B.v.S is supported by American Heart Association Postdoctoral Fellowship (#899270). A.P. is supported by NIH (K08 HL157700), Sarnoff Cardiovascular Research Foundation and Michael Antonov Charitable Foundation.

## Author Contributions

T.N. and D.S. conceived and directed the study. T.N. and Y.H. performed animal work. T.N. and S.S.R. collected heart tissues and isolated single cells for subsequent scRNA-seq and scATAC-seq. T.N., S.S.R., A.P., B.v.S, and F.K. analyzed scRNA-seq and scATAT-seq and developed computational methods. T.N. and B.v.S. performed RNA *in situ* hybridization and subsequent tissue clearing and imaging. T.N., L.Y., N.S., and M.W.C designed, performed and analyzed luciferase assay. T.N., S.S.R., M.A., A.P., J.V.B, C.A.G., M.W.C., and D.S. interpreted the data. R.T. reviewed statistical methods. T.N, M.W.C. and D.S. wrote the manuscript with contributions of M.A.

## Additional Information

Four supplementary videos are provided.

Three supplementary tables are provided.

Corresponding author: Deepak Srivastava

