## Supplemental Figures for "Single Cell Multimodal Analyses Reveal Epigenomic and Transcriptomic Basis for Birth Defects in Maternal Diabetes"

**SI Guide**

**Extended Data Tables**

**Extended Data Table 1 | The detected cardiac phenotypes at E18.5 or P0 from VEH or STZ treated females.** This table shows the number of detected cardiac phenotypes, including atrial septum defect (ASD), ventricular septum defect (VSD), atrioventricular septum defect (AVSD), and thinner right ventricle (RV) wall, at E18.5 embryos or P0 pups from VEH or STZ treated females. Representative histological images were presented in **Extended Data Figure 1b**.

**Extended Data Table 2 | Statistical results for the ChromVAR analysis.** This table shows the statistical results to determine the differences between the bias-corrected deviations for a TF motif between VEH and STZ conditions per each cell type displayed **in Extended Data Figure 8a** and **8b**. Gray highlighted cell types show the statistical differences.

**Extended Data Table 3 | Primer lists.** This table lists the used primers for cloning the candidate distal regulatory regions discussed in **Figure 4 a-d** (**Table 3a**)**.** This table lists the used primers for generating the estimated TF binding sites deletion mutants discussed in **Figure 4 a-d** (**Table 3b**)**.**

**Extended Data Movies**

**Extended Data Movie 1 | *Alx3* positive cells distribution in Mef2c-AHF-Cre; Ai6 fetus at E10.5.** Serial coronal optical sections from ventral to dorsal of whole-mount RNA *in situ* hybridization. *Alx3* (red), ZsGreen (green), and DAPI (blue) are shown. The scale bar represents 100 μm.

**Extended Data Movie 2 │ The spatial relationship between *Alx3* positive cells and *Armh4* positive cells.** The whole-mount RNA *in situ* hybridization for *Alx3* (red), *Armh4* (green) with DAPI (blue). The scale bar represents 300 μm.

**Extended Data Movie 3 │ RARE activity is enhanced in the second heart field at E10.5 in maternal diabetes. a,** The whole-mount RNA *in situ* hybridization for *Alx3* (green), LacZ (red) with DAPI (blue) in an E10.5 RARE-LacZ fetus from VEH treated female. The scale bar represents 300 μm. **b,** The whole-mount RNA *in situ* hybridization for *Alx3* (green), LacZ (red) with DAPI (blue) in an E10.5 RARE-LacZ fetus from STZ treated female. The scale bar represents 300 μm.

### Extended Data Figure Legends

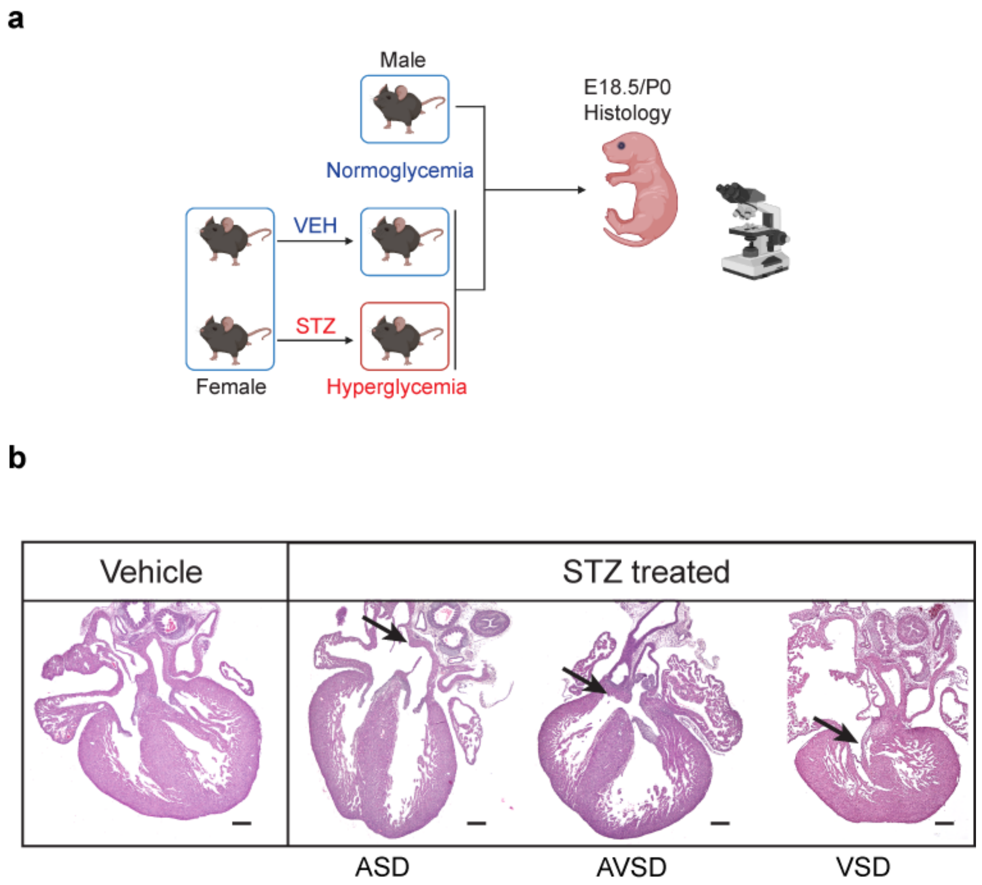


**Extended Data Fig. 1 | Histological validation of the maternal diabetes model. a,** The design of the *in vivo* maternal diabetic model experiment. After administration of either VEH or STZ, females in the STZ group with confirmed diabetes were mated with normoglycemic males, and heart samples at embryonic day 18.5 (E18.5) or postnatal day 0 (P0) were collected for histological examination. **b,** Representative images of the heart phenotypes detected in the diabetic model. The prevalence of each malformation is shown in Extended Data Table 1. The scale bar represents 200 μm.

**Extended Figure 2**

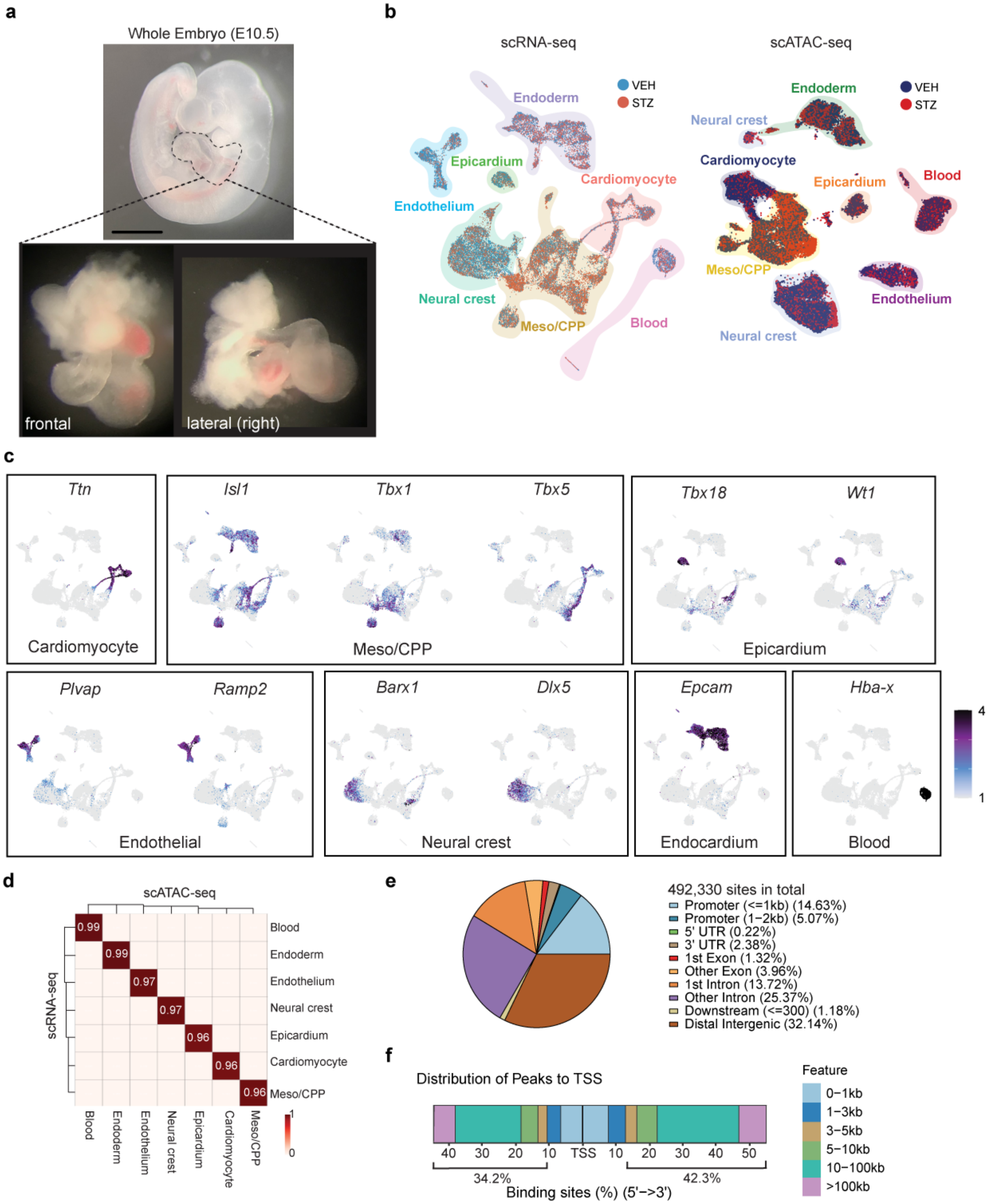


**Extended Data Fig. 2 | Single Cell Multimodal Analysis of Cardio-pharyngeal Region in Maternal Diabetes. a,** Representative image of E10.5 embryo with detailed micro-dissected region used for scRNA/scATAC-seq experiment. Scale bar represents 1 mm. **b,** scRNA-seq (left) or scATAC-seq (right) UMAP presentation colored by conditions. Different colors overlayed delineate cell type cluster annotations. **c,** Expression patterns of representative cell type specific marker genes plotted on UMAP space shown in Figure 1b. **d,** Heatmap of Jaccard indices calculated between scRNA-seq and scATAC-seq after integration. Values range from 0 to 1 (higher value represents closer annotation matching between the two modalities). **e,** Genomic distribution of all the called peaks color coded by the genomic location as shown. Total called peaks = 492,330. **f,** Distribution of all called peaks based on the distance from transcription start sites.

### Extended Figure 3

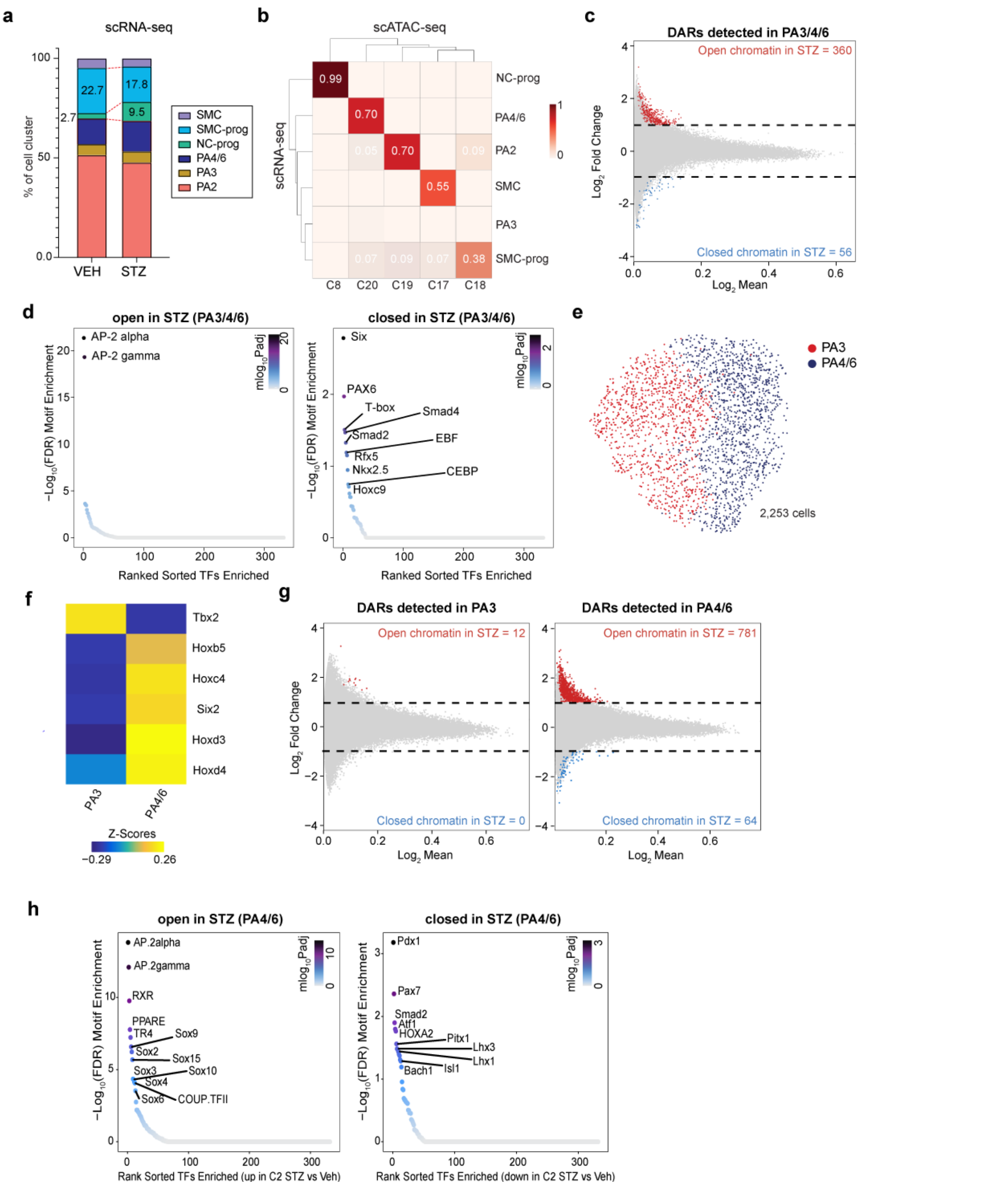


**Extended Data Figure 3 | Maternal Diabetes Dysregulates Epigenomic Landscape of Pharyngeal Arches 4 and 6 Neural Crest Cells. a,** Population distribution by sub-cell-type normalized to total number of cells per sample in neural crest cell subset data of scRNA-seq. Numbers inside the barplot represent the percentage of cell types of the total cell number. Statistics performed by permutation test in scRNA-seq data, comparing STZ vs. VEH, for NC-prog, FDR<0.001, Log2FD=1.88; for SMC-prog, FDR<0.001, Log2FD=-0.33. **b,** Heatmap of Jaccard indices calculated between neural crest cell scRNA-seq and scATAC-seq cell annotations after integration. Values range from 0 to 1 (the higher value represents closer annotation matching between those two modalities). **c,** MA plot of DARs in PA3/4/6 population between VEH and STZ. Red dots represent the more accessible (open) (FDR <= 0.05 & Log2FC >= 1) and blue dots represent less accessible (closed) DARs in STZ (FDR <= 0.05 & Log2FC <= -1). **d,** Enriched TF binding motifs in more accessible (left) or less accessible (right) DARs in STZ vs. VEH within the PA3/4/6 population. **e,** scATAC-seq UMAP representation of neural crest cell C20 subset population colored by clusters (PA3 – dark red; PA4/6 – dark blue). **f,** Heatmap of Gene Scores (GS) of curated marker genes based on scRNA-seq data for PA3 and PA4/6 neural crest. Scale indicates z-scored GS values. **g,** MA plot of DARs between VEH and STZ in PA3 population (left) and PA4/6 population (right). Red dots represent the more accessible (open) (FDR <= 0.05 & Log2FC >= 1) and blue dots represent less accessible (closed) DARs in STZ (FDR <= 0.05 & Log2FC <= -1). **h,** Enriched TF binding motifs in more accessible (left) and less accessible (right) DARs in STZ in PA4/6 population. NC-prog, neural crest cell progenitors; PA2, pharyngeal arch 2; PA3, pharyngeal arch 3; PA4/6, pharyngeal arch 4/6; SMC, smooth muscle cells; SMC-prog, smooth muscle cell progenitors.

### Extended Figure 4

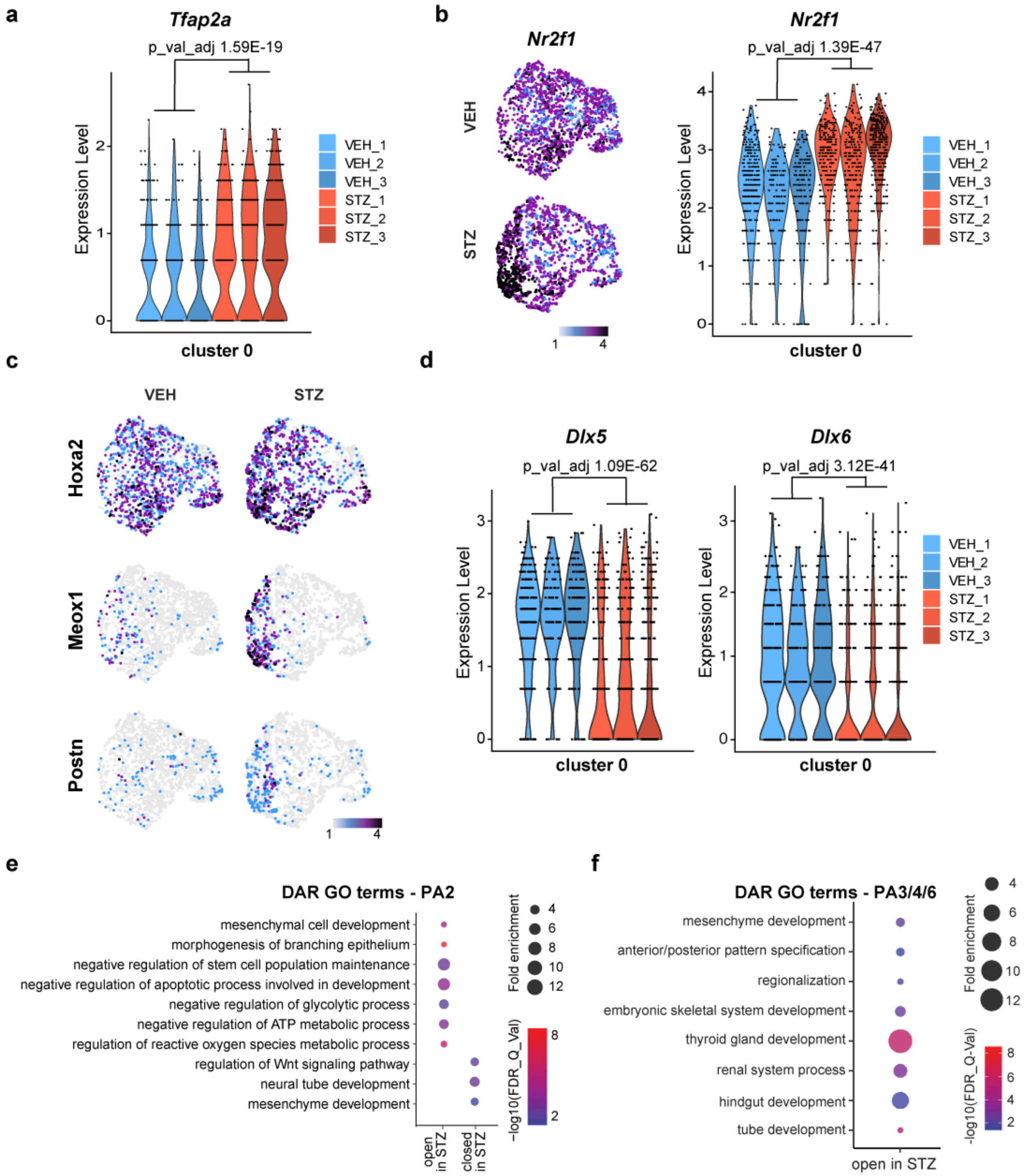


**Extended Data Fig. 4 │ Maternal Diabetes Dysregulates Epigenomic and Transcriptional Landscape Associated with Cell Differentiation and Patterning in Pharyngeal Arch Neural Crest. a,** Violin plot of *Tfap2a* expression levels in cluster 0 of

UMAP in Figure 2i across 3 VEH and 3 STZ embryos. **b,** Expression of *Nr2f1* mRNA on UMAP space for PA2 neural crest cells (VEH – left top; STZ – left bottom). Scale bar indicates z-scored expression values. Violin plot of *Nr2f1* expression levels in cluster 0 of UMAP in Figure 2i (right). **c,** Expression of indicated genes on UMAP space for PA2 neural crest cells. Scale bar indicates z-scored expression values. **d,** Violin plots of *Dlx5* (left) and *Dlx6* (right) expression levels in cluster 0 of UMAP in Figure 2i. **e,** Enriched GO terms in detected DARs in PA2 population using GREAT analysis. **f,** Enriched GO terms in detected DARs in PA3/4/6 population using GREAT analysis.

### Extended Figure 5

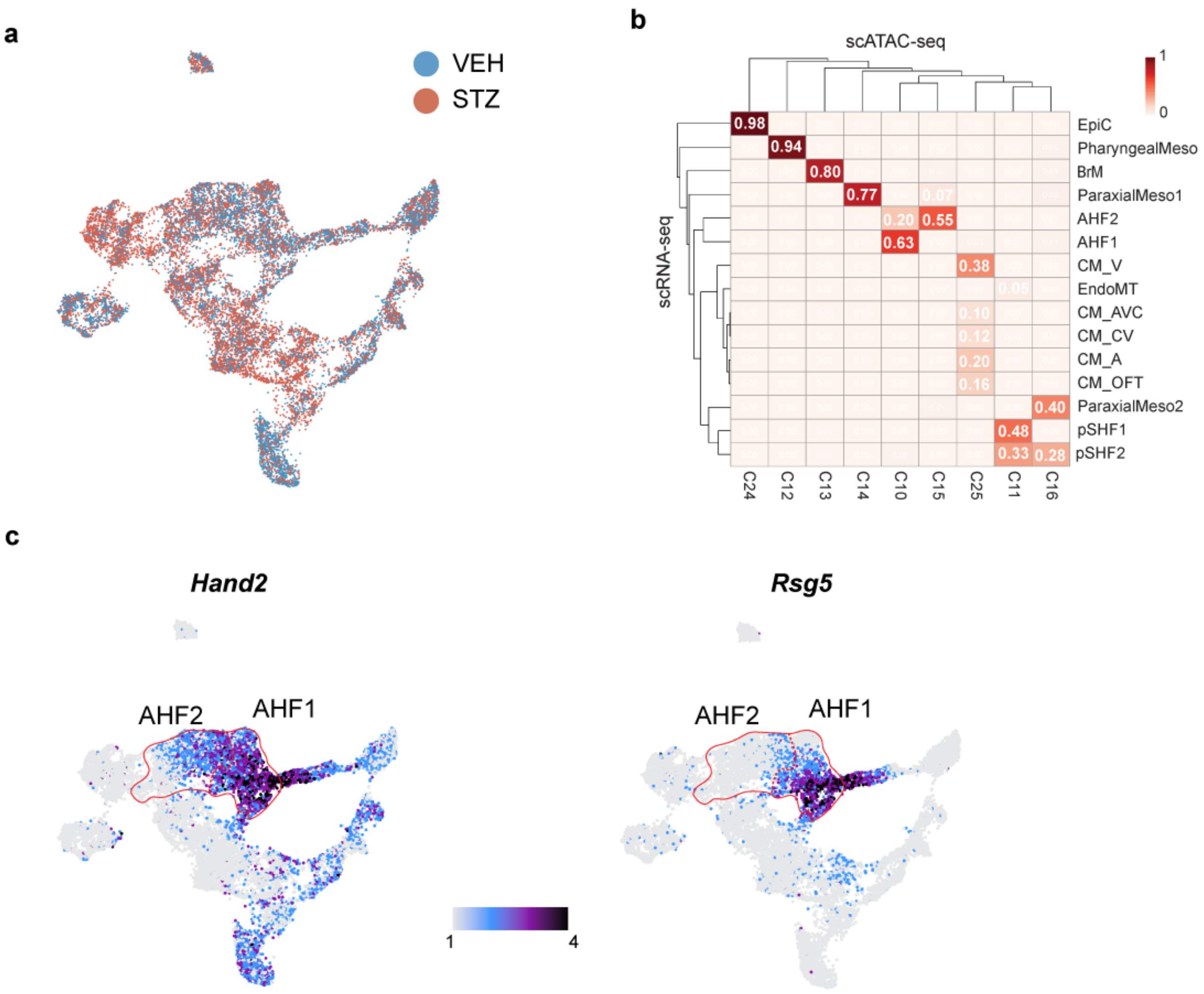


**Extended Data Fig. 5 | Maternal Diabetes Specifically Affects *Alx3*$^{pos}$ AHF cells, Causing Abnormal Anteroposterior Patterning. a,** scRNA-seq UMAP representation of mesodermal population (Meso/CPP", "Cardiomyocyte", or "Epicardium" in Figure 1b) colored by conditions (VEH – blue; STZ – light red). **b,** Heatmap of Jaccard indices between mesoderm cell scRNA-seq and scATAC-seq annotations after integration. Values range from 0 to 1 (the higher value represents closer annotation matching between those two modalities). CM_V, ventricular cardiomyocyte; CM_AVC, atrioventricular canal cardiomyocyte; CM_A, atrial cardiomyocyte; CM_SV, sinus venosus cardiomyocyte; CM_OFT, outflow tract cardiomyocyte; pSHF1/2, posterior second heart field 1/2; EndoMT, endothelial mesenchymal transition; EpiC, Epicardium; AHF1/2, anterior heart field 1/2; PharyngealMeso, pharyngeal mesoderm; ParaxialMeso1/2, paraxial mesoderm 1/2; BrM, branchiomeric muscle. **c,** Expression pattern of *Hand2* (left) and *Rgs5 (right)* on UMAP space. AHF1 and 2 are circled in red. Scale bar indicates z-scored expression values.

### Extended Figure 6

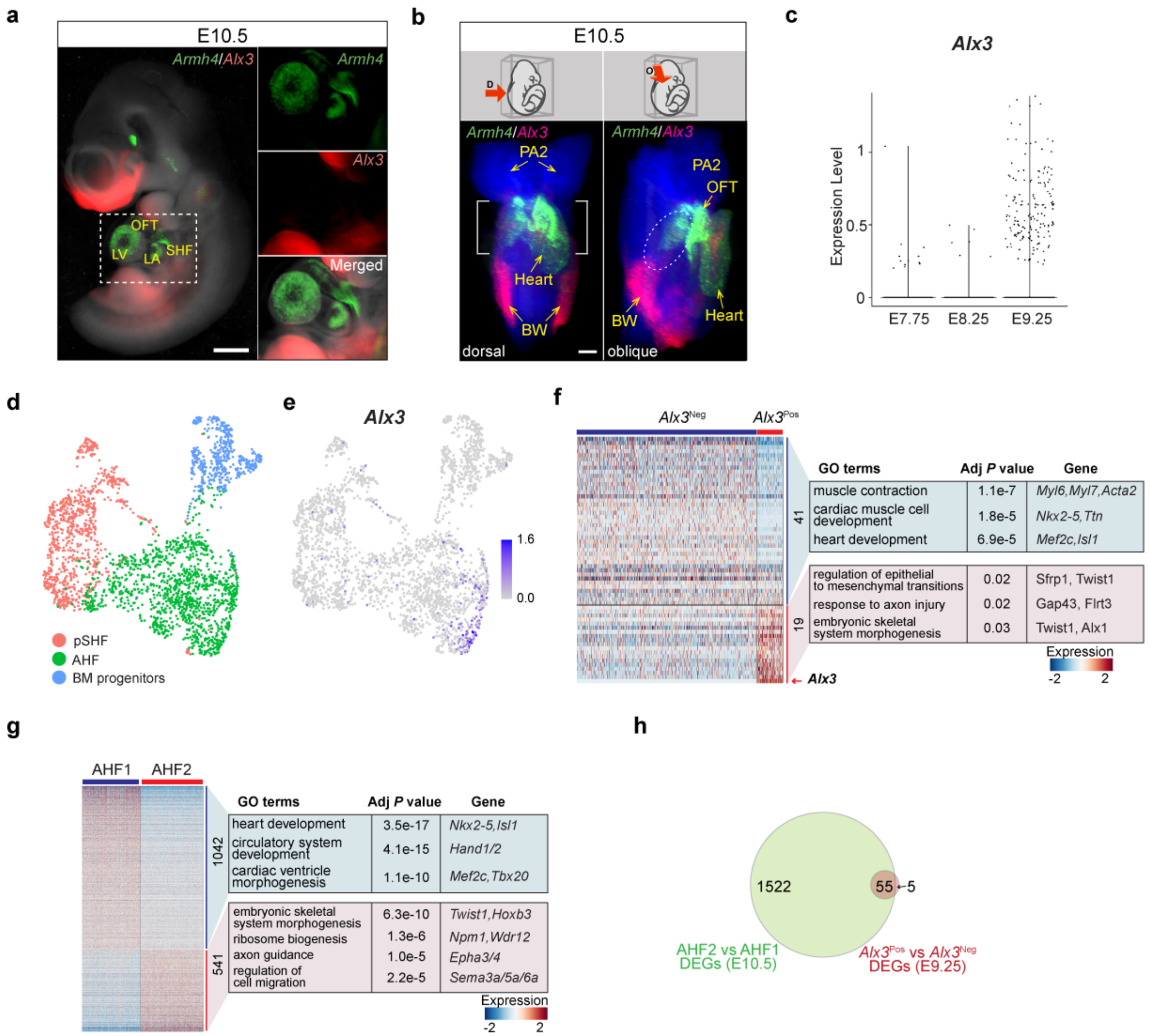


**Extended Data Fig. 6 │ *Alx3*$^{Pos}$ cells are a Distinct Subset of the AHF Population.**

**a,** Representative images form RNA *in situ* hybridization for *Armh4* (green) and *Alx3* (red) in an E10.5 embryo from VEH treated female. The scale bar represents 500 μm. **b,** Representative images from whole mount RNA *in situ* hybridization of E10.5 embryos using light sheet microscopy. *Armh4* (green) and *Alx3* (red) expression is shown from the dorsal view (D – left) and the right oblique view (O – right). A white bracket (left) highlights the anterior part of *Alx3*$^{Pos}$ cells. A white dotted oval (right) highlights the

*Alx3*$^{\mathrm{Pos}}$ cell streak on left side of the embryo from outflow tract (OFT) towards the posterolateral region. Still images were extracted from Extended Data Movie 2. Scale bar represents 100 μm. PA2, pharyngeal arch 2; BW, body wall. **c,** The distribution of *Alx3* positive cells by scRNA-seq between E7.75 and E9.25. **d,** scRNA-seq UMAP of cardiac progenitor cells at E9.25 from the same data as **(c)** color coded by cell type annotation. AHF, anterior heart field; BM progenitors, branchiomeric muscle progenitors; pSHF, posterior second heart field. **e,** Expression of *Alx3* on the same UMAP as **(d)**. **f,** Heatmap of differentially expressed genes (DEGs) between *Alx3*$^{\mathrm{Neg}}$ AHF and *Alx3*$^{\mathrm{Pos}}$ AHF at E9.25. All detected DEGs that attained adjusted p-val < 0.05 and Log$_2$FC > 0.25 are shown. Top GO terms enriched in upregulated or downregulated DEGs are shown with representative genes composing each GO. Scale bar indicates z-scored expression values. **g,** Heatmap presentation of DEGs between AHF1 and AHF2 using only VEH cells in the scRNA-seq data. All detected DEGs that attained adjusted p-val < 0.05 and Log$_2$FC > 0.25 are shown. Top GO terms enriched in upregulated or downregulated DEGs are shown with representative genes composing each GO. Scale bar indicates z-scored expression values. **h,** Venn diagram representing the intersect between DEGs shown in **(f)** and **(g)**.

### Extended Figure 7

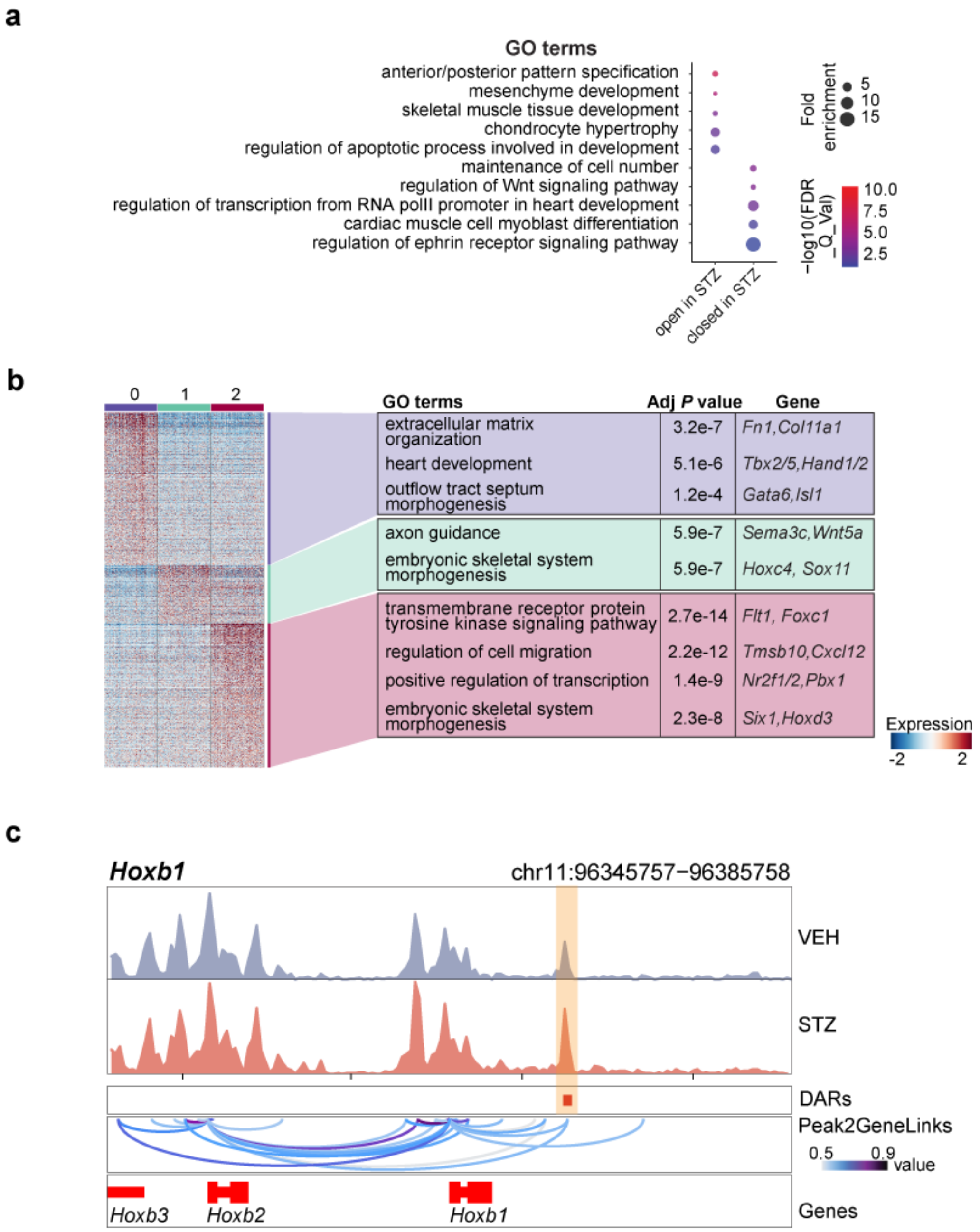


**Extended Data Fig. 7 | PGDM Disrupts Anterior-Posterior Patterning in AHF2. a,** Enriched GO terms in VEH vs. STZ DARs in AHF2 population using GREAT analysis. **b,** Heatmap of marker genes of each of three subclusters found in *Alx3*$^{Pos}$ AHF2. These marker genes were detected using only VEH-treated *Alx3*$^{Pos}$ AHF2 cells (left). All marker genes that attained an adjusted p-val < 0.05 and $Log_2FC > 0.25$ are shown. Scale bar indicates z-scored expression values. Top GO terms enriched in marker genes for each sub cluster with statistical information and representative maker genes to corresponding GO term are shown (right). **c,** Genome browser plots for *Hoxb1* locus. The top two rows represent the chromatin accessibility in VEH and in STZ within AHF2. The third track from the top shows the genomic location of the DAR with more accessibility in STZ (red rectangles, highlighted by yellow box). The second track from the bottom represent the links between peaks and gene ("Peak2GeneLinks"), calculated by ArchR. Darker lines represent stronger links. The bottom track shows the gene location and transcriptional direction (red – positive strand; blue – negative strand).

### Extended Figure 8

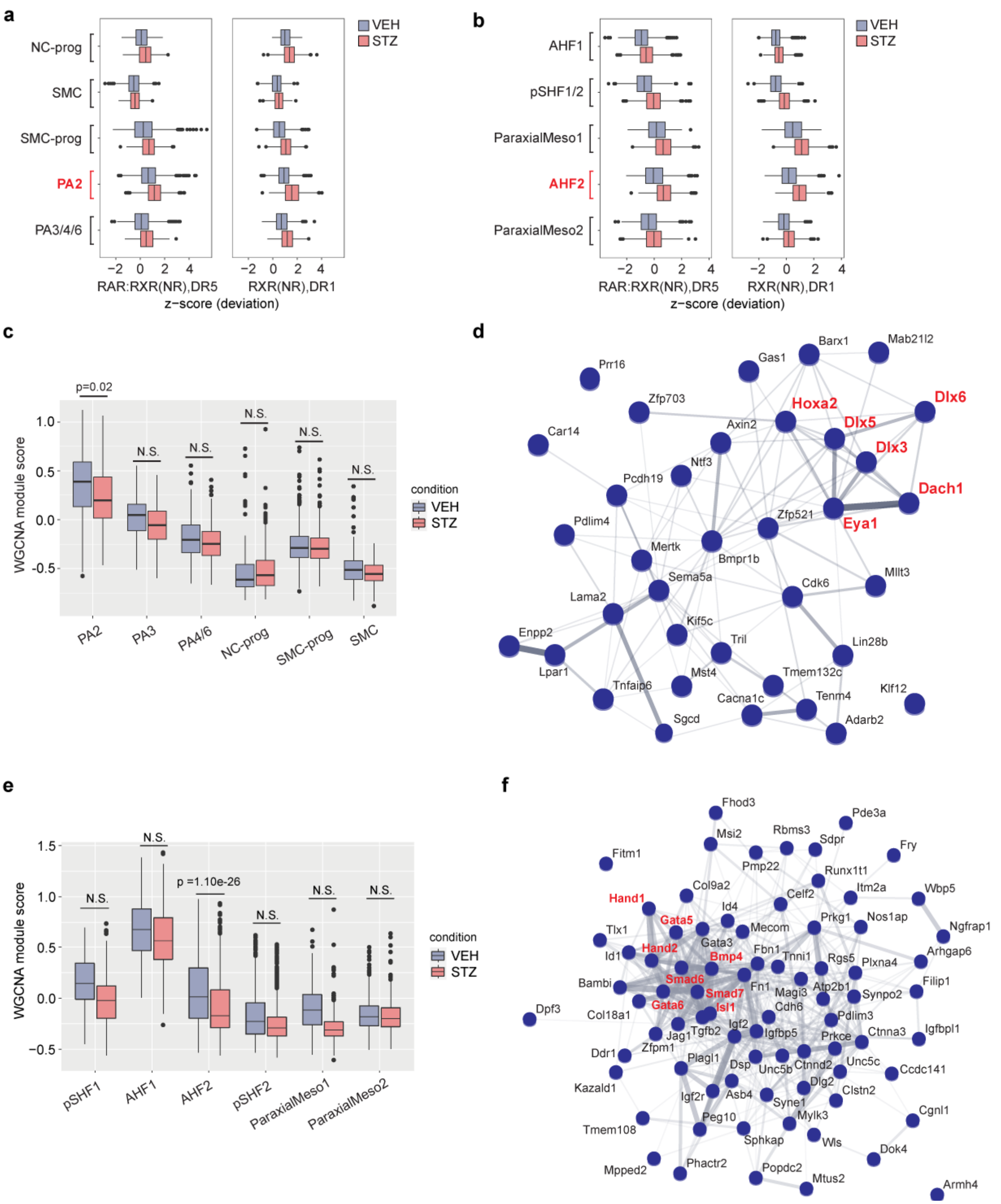


**Extended Data Fig. 8 | Disrupted Retinoic Acid Signaling is Associated with Dysregulation of Gene Regulatory Networks in Pharyngeal Arch 2 and AHF2. a,** Box plots of the distribution of ChromVAR deviation score for RAR and RXR transcription factor motifs for each cluster in the neural crest cell population. PA2 neural crest cells are highlighted in red. The X-axis shows the distribution of the Z-score. (STZ – red; VEH – blue). **b,** Box plots of the distribution of ChromVAR deviation score for RAR and RXR transcription factor motifs for each cluster in the mesoderm population. AHF2 cells are highlighted in red. The X-axis shows the distribution of the Z-score. (STZ – red; VEH – blue). **c,** Module scores for a gene module detected in the WGCNA analysis that showed statistically significant variation between VEH and STZ only in PA2. Linear mixed effects models with mouse id as the random effect was used to test the significance of the mean difference in the module score between VEH and STZ. **d,** Map of functional protein-protein interactions (PPI) of genes composing the module described in c, depicted using STRING. Genes composing a core of the PPI network and being downstream of *Tfap2* are highlighted in red and bold. **e,** Module scores for a cardiac gene regulatory module detected in the WGCNA analysis that showed statistically significant variation between VEH and STZ only in AHF2. The same statistical test as **(c)** was used. **f,** Map of PPI of genes composing the module described in d, depicted using STRING. Genes composing a core of the PPI network and being critical cardiac TFs or signaling genes are highlighted in red and bold. NC-prog, neural crest cell progenitors; PA2, pharyngeal arch 2; PA3, pharyngeal arch 3; PA4/6, pharyngeal arch 4/6; SMCs, smooth muscle cells; SMC-Prog, smooth muscle cell progenitors; pSHF1/2, posterior second heart field 1/2; AHF1/2, anterior heart field 1/2; ParaxialMeso1/2, paraxial mesoderm 1/2.
